# Non Hybrid Long Read Consensus Using Local De Bruijn Graph Assembly

**DOI:** 10.1101/106252

**Authors:** German Tischler, Eugene W. Myers

## Abstract

While second generation sequencing led to a vast increase in sequenced data, the shorter reads which came with it made assembly a much harder task and for some regions impossible with only short read data. This changed again with the advent of third generation long read sequencers. The length of the long reads allows a much better resolution of repetitive regions, their high error rate however is a major challenge. Using the data successfully requires to remove most of the sequencing errors. The first hybrid correction methods used low noise second generation data to correct third generation data, but this approach has issues when it is unclear where to place the short reads due to repeats and also because second generation sequencers fail to sequence some regions which third generation sequencers work on. Later non hybrid methods appeared. We present a new method for non hybrid long read error correction based on De Bruijn graph assembly of short windows of long reads with subsequent combination of these correct windows to corrected long reads. Our experiments show that this method yields a better correction than other state of the art non hybrid correction approaches.

## 1 Introduction

First generation sequencing allowed to determine the genomic sequences of important genomes like the human (cf. [14, 28]) and fly (see [3]) genomes. While the technology was suitable for achieving these very important goals, it was too expensive and slow for many applications. Second generation sequencing brought the advent of high throughput sequencing and was much more affordable and quick. Second generation technologies however have, especially with the application of genome assembly in mind, the major drawback of producing much shorter reads than those which were common in the first generation. Second generation reads, which are often no more than 150 base pairs (bp) long, enable a much lower capability to correctly resolve genomic repeat regions than the previously used first generation reads with an average read length of 700 bp. Third generation sequencers like those built by Pacific Biosciences (PacBIO) and Oxford Nanopore yield much longer reads up to 50000 bp with an average of 15kb for PacBIO. In addition these sequencers can work on single molecules, which in principle makes polymerase chain reaction (PCR) unnecessary and thus removes the bias induced by this procedure. These features however come at the price of a greatly enlarged average base error rate of 15% and higher. This poses enormous algorithmical challenges. To work effectively with the data obtained in many cases, including single nucleotide polymorphism (SNP) detection and genome assembly, requires correcting most of these errors. Second generation error correction handled mainly the substitution errors typical for short read data and is not suitable to handle long read data in which most errors are insertions or deletions. The first algorithmic approaches for correcting errors in third generation reads used second generation reads (see e.g. [4, 10, 18, 24]). These approaches are called hybrid as they combine two different types of sequencing data. Such approaches however inevitably suffer from two problems. First it is often unclear where to map the short read data on the long reads if long reads span repeats which cannot be resolved by short reads. Secondly the short read data suffers from amplification biases so for certain regions which can be successfully sequenced using third generation technology there will be no coverage by second generation data. More recently non hybrid methods based on third generation data only were established (see for instance [6, 7, 12, 25]). In this paper we present a new method for non-hybrid long read error correction based on local De Bruijn graph assembly. While genome wide De Bruijn graph based assembly is infeasible using long read data, we show that even for high error rates the De Bruijn graph approach is effective on small windows. Experiments show our approach to be competitive with previously published work.

## 2 Method

### 2.1 Modelling Sequencing Events

For the sake of modelling let us assume that in comparison with the true sequence a sequencer may produce three kinds of error events:

- insertion (I): the sequencer reports a base which has no corresponding base in the true sequence
- deletion (D): the sequencer fails to report a base existing in the true sequence
- substitution (S): the sequencer reports a base which differs from the one found in the true sequence and a single type of event M representing a correctly reported base. Let X_i_ for *i* = 0, 1, … denote a sequence of independent and identically distributed (iid) discrete random variables such that 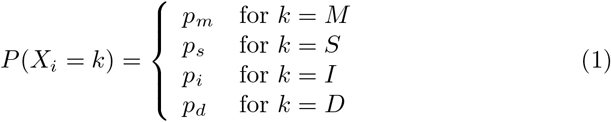

and *p*_*m*_ + *p*_*s*_ + *p*_*i*_ + *p*_*d*_ = 1. The sequence X_*i*_ represents the series of events produced by a sequencer. In this setting the probability to see *n*_*i*_ insertions before the *j*’th non insertion event is given by a negative binomial distribution, the probability mass function (PMF) of which is given by 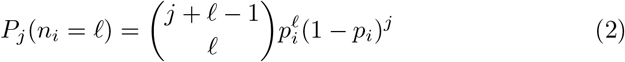

The probability to see *n*_*d*_ deletions within the first *j* non insertion events follows a binomial distribution. It’s PMF is

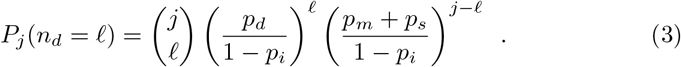

Given these two functions we can deduce the probability to see a correctly reported base at true position *t* offset by *o* in a sequenced read (i.e. the base is at position *t* in the true sequence but *t* + *o* in a sequenced read) as

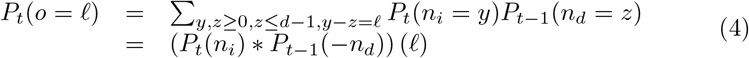

where the * symbol denotes the convolution operator.

Figure 1 shows *P*_*t*_ for several values of *t* and error probabilities *p*_*i*_, *p*_*d*_ and *p*_*s*_ typically found in PacBIO reads.

**Figure 1:**
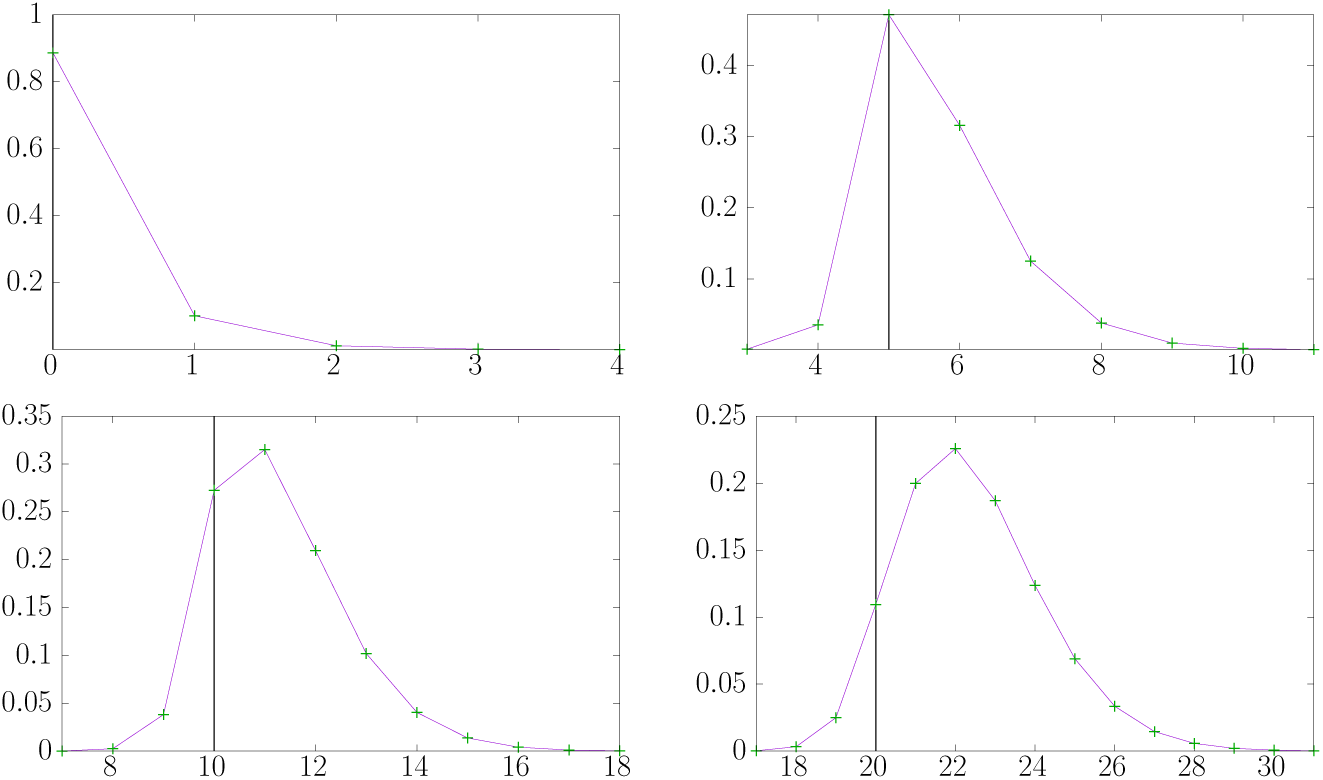
Function *P*_*t*_(*o*–*t*) for *t* = 0 (top left), *t* = 5 (top right), *t* = 10 (bottom left) and *t* = 20 (bottom right) using *p*_*i*_ = 0:114, *p*_*d*_ = 0:0135 and *p*_*s*_ = 0:0121 implying an error rate of 14%. The horizontal axis denotes positions in the sequenced read. The vertical axis represents the probability of seeing a correctly reported base from position *t* in the true sequence at the respective *x* position in a read. The vertical black bar marks the respective value of *t* in each graph. Note that for *t* = 10 and *t* = 20 the maximal probability does not appear at *x* = 10 and *x* = 20 respectively as insertions are far more likely than deletions.

### 2.2 Edit Scripts

Let ∑ = {*A,C, G,T*} denote the DNA alphabet. The set OP = {*M*; *S*_*A*_; *S*_*C*_; *S*_*G*_; *S*_*T*_; *I*_*A*_; *I*_*C*_; *I*_*G*_; *I*_*T*_; *D*} contains the edit operations we consider (copy/match, substitute/replace by *A, C, G, T*, insert *A, C, G, T* and delete). An edit script *S* is a sequence over OP. For an edit script *S* = *s*_1_, *s*_2_, … *s*_i_ the occurrence count function Occ_*O*_ defined by 
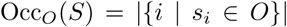
 for any subset *O* ⊆ OP counts the number of operations from *O* contained in *S*. The number of errors in an edit script is given by the err function defined by err(*S*) = Occ_OP\{*M*}_(*S*) and the error or difference rate function erate for a script *S* by

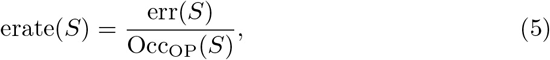

i.e. the number of error operations in *S* divided by the number of all operations in *S*. An edit script *S* is admissible for a string *R* of length |*R*| over ∑ if Occ_{*M*, *S*_*A*_, *S*_*C*_, *S*_*G*_, *S*_*T*_, *D*}_(*S*) = |*R*|, i.e. the number of non-insertion operations in *S* equals the number of symbols in *R*. Given a string or sequence *S* = *s*_1_, …, *s*_n_ and *i*, *j* s.t. *i* ≤ *j* we define *S*_i.j_ by *S*_i_*S*_i+1_ … *S*_j_ if 1 ≤ *i* ≤ *j* ≤ *n* and the empty string/sequence respectively otherwise. We use *S*_i_ as a short version of *S*_i,i_. For a string *R* over and an edit script *S* admissible for *R* the edit function edit (*R*, *S*) is recursively defined as

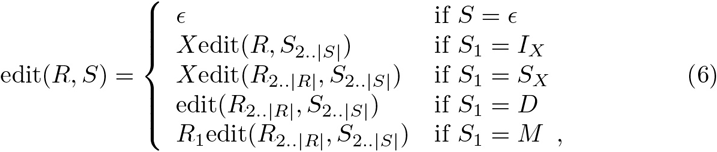

i.e. both *R* and *S* are scanned from left to right and the edit operations in *R* are applied to *S*. Given two strings *U* and *V* over ∑ and optimal alignment of *U* and *V* is any edit script *S* admissible for *U* s.t. edit(*U*, *S*) = *V* which minimises the err function.

### 2.3 Read Alignment Piles

Let *G* = {*G*_1_, *G*_2_,…, *G*_g_denote a genome, i.e. each *G*_i_ is a string over the DNA alphabet ∑ = {*A,C,G,T*}. Further, let *P* = *P*_1_,…, *P*_p_ denote a set of read parameters s.t. each *P*_i_ is a tuple (*j*; *b*; *e*; *d*; *S*) where 1 ≤ *j* ≤ *g*, 1 ≤ *b* ≤ *e* ≤ |*G*_*j*_|, *d* ∈ {0, 1} (forward or reverse complement) and *S* is an edit script admissible for *G*_b,e_. We denote the components of any such tuple using the notation *P*_i_. *j*, *P*_i_. *b*, etc. Each element of *P* denotes a sequencing read. It specifies where in the genome the read stems from (coordinates and strand) and how this region needs to be modified to obtain a resulting read. More precisely a read is obtained from a read parameter tuple by extracting *G*_j__b,e_, applying the reverse complement operation if *d* is set and finally applying the edit script *S*. We denote the read described by a tuple *P*_i_ as read(*P*_i_). Two reads given by the tuples *P*_i_ and *P*_k_ overlap in *G* if *P*_i_. *j* = *P*_k_. *j* and [*P*_*i*_. *b*, *P*_*i*_. *e*] ∩ [*P*_*j*_. *b*, *P*_*j*_. *e*] ≠ ∅. Given two overlapping reads by *P*_*i*_ and *P*_*k*_ we can deduce an edit script representing their relative alignment on G (details are described in Appendix A). Aligning read(*P*_*i*_) and read(*P*_*k*_) using the edit script obtained requires applying the reverse complement operations as stored in *P*_*i*_. *d* and *P*_*k*_. *d* respectively. In general any alignment of two reads *A* and *B* can be described as an alignment tuple (abpos; aepos; bbpos; bepos; inv; *S*) where abpos denotes the start position of the alignment on *A*, aepos the end position on *A*, bbpos and bepos start and end position on *B* respectively, inv whether the *B* read is to be reverse complemented and *S* the edit script, which needs to be admissible for *A*_abpos;aepos_ and satisfy edit(*A*_abpos;aepos_; *S*) = *B*_bbpos,bepos_ if inv = 0 and edit 
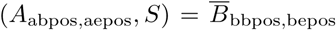
 otherwise (we use 
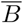
 to denote the reverse complement of *B*). One easily checks that it is sufficient to have the inverse complement option for one of the reads only. If we would need to invert both reads instead, then this would be equivalent to keeping the reads as they are and reversing the edit script instead. For a given read *A* we call any set of alignment tuples between *A* and other reads an alignment pile for *A*. Note that in this definition we do not require that *A* truly overlaps the reads contained in the pile in the sense that both *A* and all the other reads stem from the same place in a genome. The definition of alignment tuples mandates the validity of the represented alignments though. This reflects the scenario we mostly find in real data. We see significant (local) alignments between reads, but there is no guarantee that these local alignments all represent true overlaps in the underlying genome. In contrast to this we define the perfect alignment pile for a read read(*P*_*i*_) given by a tuple *P*_*i*_ as the set of alignment tuples constructed from all reads *P*_*j*_ overlapping *P*_*i*_ on *G*. Figure 2 depicts a perfect alignment pile for a read described by some tuple *P*_*i*_.

**Figure 2:**
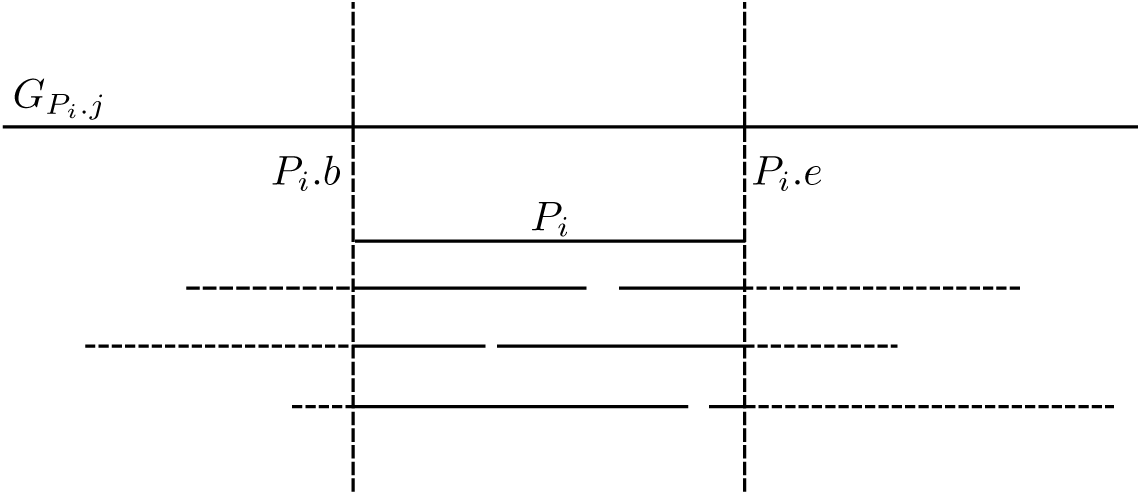
Perfect alignment pile for a read described by a tuple *P*_*i*_. Other reads overlapping *P*_*i*_ are drawn solid inside the pile and dashed outside the pile.

### 2.4 De Bruijn Graph

The De Bruijn graph is a data structure used in many assembly programs (see e.g. [23, 26, 17]). It was first introduced as a concept for genome assembly in [11]. Let ∑ = {*A,C, G, T*} denote the DNA alphabet and let *R* = {*R*_1_, *R*_2_,…, *R*_r_} denote a set of strings (reads) over ∑. The De Bruijn graph *D* = (*V*, *E*) of order 
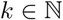
 *R* is given by

- the set of vertices *V* containing exactly the substrings of length k (so called *k*-mers) occurring in the strings in *R* and
- the set of edges *E*, where an edge exists from *v*_1_ = a_1_a_2_: a_k_ to *v*_2_ = b_1_b_2_: b_k_ i a_2_a_3_: a_*k*_ = b_1_b_2_: b_*k*1_, i.e. *v*_1_ and *v*_2_ overlap by *k*−1 symbols/bases.

The essential idea used for assembly is that, given a set of preconditions, the genome is contained in the graph as a set of paths/tours through the graph. The necessary condition, which we call connectivity, for this is that each *k*-mer appearing in the genome appears in at least one read. Assuming the reads represent a uniform sampling of the genome at coverage *c* (i.e. each base is represented in c reads) the probability P_*k*_ to see this happen for a single k-mer is bounded below by 1 − (1−*p*^*k*^)^*c*^ if the behaviour of the sequencer can be formulated as an iid time discrete stochastic process with probability *p* to see a correctly reported base. This is a lower bound as the *k*-mer could appear more than once in the genome. This function converges to 1 for increasing values of *c* if *p* ≠ 0. However, depending on the size of *k* it may require an infeasibly high coverage *c* to see it happen with a probability close to 1. In particular the higher *k* is, the lower *P*_*k*_ gets for constant *c*. For this reason a lower value of *k* makes it more likely to fulfil the necessary condition to allow a set of paths through the graph spelling out the sequences of the genome. The left plot in Figure 3 shows the probability to see a *k*-mer with increasing depth for various values of *k*.

**Figure 3:**
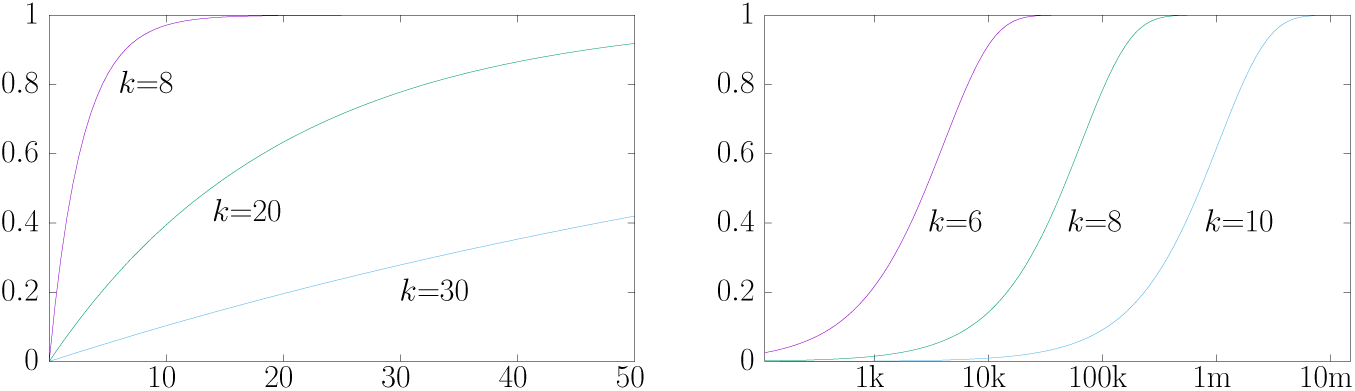
Left: probability to see a correct *k*-mer (*k* = 8; 20 and 30) in a set of reads of coverage *c* with a single base error rate of 14%, the horizontal axis denotes *c* and the vertical the probability. Right: probability to see another copy of a given *k*-mer (*k* = 6; 8 and 10) in a genome, the horizontal axis denotes the length of the genome observed, the vertical marks the probability

On the other hand a lower value of *k* makes it more likely to get a graph with a lot of branching nodes, i.e. such nodes which have more than one successor. Each of these makes it computationally harder and practically more unlikely to find paths in the graph which represent correctly reconstructed sequences from the genome. For this reason a higher value of *k* would be bene cial. In a genome which has size *G* ≥ 4^*k*^ (the length of the corresponding De Bruijn sequence) and where each *k*-mer appears with the same probability the likelihood of seeing certain *k*-mer is

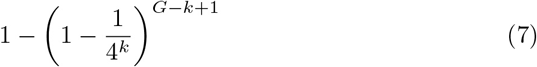

Which is also roughly the likelihood of seeing any given *k*-mer again in a genome of size *G* after having observed it once. The plot on the right side of Figure 3 shows this function for several values of *k*. Any such repeating *k*-mer node has a probability of 
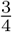
 to be a branching node.

The probability *p* of a correct base is a feature of the sequencing technology employed and in most cases cannot be changed easily. The sequencing coverage feasible is constrained as well in practice, mostly so by financial considerations.

The term contig denotes a contiguously reconstructed stretch of a genome from read data. For a set *S* = {*C*_1_, *C*_2_,…, *C*_*n*_} of contigs such that w.l.o.g. |*C*_1_| ≥ |*C*_2_| ≥ … ≥ |*C*_*n*_| the n50 statistic n50(*S*) is |*C*_*j*_| for the smallest *j* s.t. 
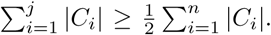

Figure 4 shows the maximum n50 contig length obtainable for the species E.coli and D.mel using De Bruijn graphs of varying orders when following non-branching paths through the graphs. In this context a branching node is any node with more than one predecessor or successor in the graph. A non-branching path is a path starting and ending in a branching node while not traversing any branching nodes in between. For a sensible assembly result one would desire that the reconstructed contigs in most cases at least exceed the average read length, which e.g. for the PacBIO sequencing machines is about 14*k* on average at the time being. The graphs show that in a perfect scenario of error free reads this happens for *k* = 24 in the case of E.coli and *k* = 30 for D.mel. Figure 3 shows that for such *k* and error rates typical for long read data a whole genome scale assembly using a standard De Bruijn graph at a coverage of 50*x* would be infeasible, as many of the *k*-mers appearing in the genome would not appear in even a single read. An approach using a sparsified version of the graph as proposed in [15] may be feasible though.

**Figure 4:**
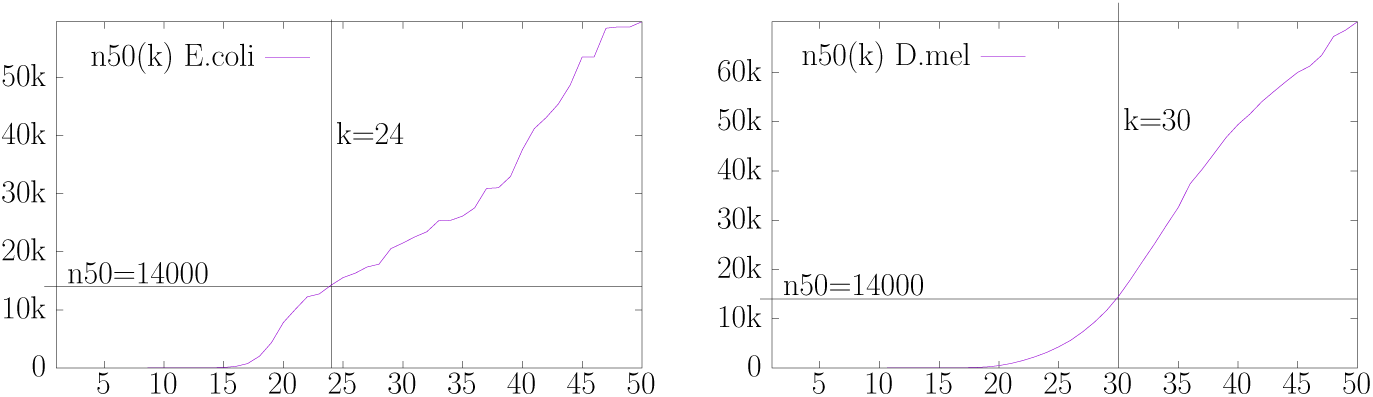
Contig n50 size obtained by following non-branching paths in De Bruijn graphs for E.coli (left) and D.mel (right). The horizontal axis shows the order of the De Bruijn graph used, the vertical axis the contig length obtained.

The situation changes notably if the scenario is switched from whole genome assembly to the assembly of small windows on a genome covering no more than 100 base pairs. Figure 5 shows a zoom of the right hand side plot depicted in Figure 3. The probability to see a second copy of a given *k*-mer is very low even for small values of *k* between 7 and 10 if we assume the window considered is produced by an iid process with equal base probabilities. The probability of a repeating *k*-mer can be substantially higher for real genomes than in such iid sources. We call a De Bruijn graph built on a perfect single read representing a window of a genome a window De Bruijn graph. Figure 6 shows that for genomes like E.coli and D.mel the window De Bruijn graphs for windows of size 40 and 100 can in most cases be traversed while encountering a very low number of branching nodes to reclaim the original window. So even for the more repetitive case of real genomes it is still feasible to perform De Bruijn graph based assembly of small windows based on noisy long read data. Note that while the number of branching nodes is low in this scenario, there is still a significant number of windows whose reconstruction requires visiting at least one branching node. We will tackle this issue in the following.

**Figure 5:**
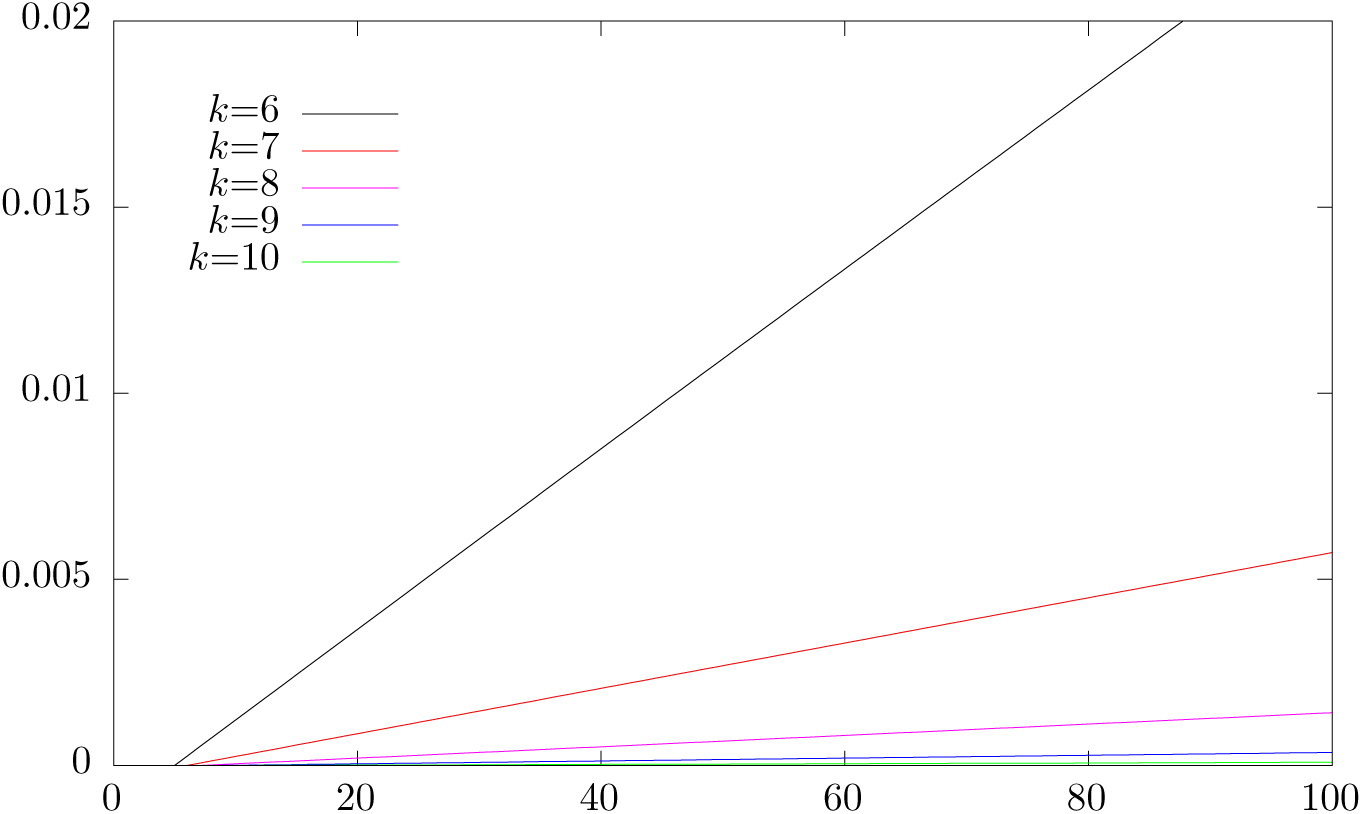
Probability to see another copy of a given *k*-mer (*k* = 6; 7; 8; 9 10) in a window of a given size. The horizontal axis denotes the window size, the vertical marks the probability.

**Figure 6:**
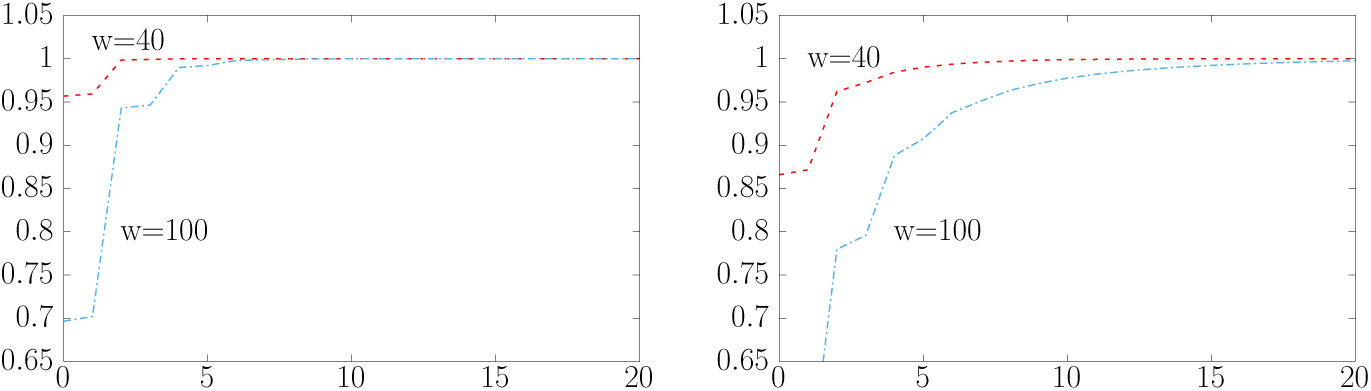
Fraction of windows of size 40 and 100 in E.coli (left) and D.mel (right) having De Bruijn graphs of order *k* = 8 with a path representing the window featuring up to a given number of branching nodes. The horizontal axis denotes the number of branching nodes and the vertical axis the probability.

### 2.5 *k*-mer scores

Let *R* = {*R*_1_, *R*_2_, …, *R*_r_} denote a set of strings/reads over the DNA alphabet and let 0 ≤ *p*_*m*_; *p*_*s*_; *p*_*i*_; *p*_*d*_ ≤ 1 sequencing event probabilities s.t. *p*_*m*_+*p*_*s*_+*p*_*i*_+*p*_*d*_ = 1. Further let 
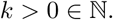
 W.l.o.g. assume |*R*_i_| ≥ *k* for all 1 ≤ *i* ≤ *r*, if any of the reads is too short, then we just drop it. Each read *R*_i_ generates a sequence of pairs of a *k*-mer and a position. For instance if *R*_*i*_ = *c*_1_*c*_2_… *c*_*n*_ for *n* ≥ *k*, then the read generates the pairs (*c*_1.k_; 0); (*c*_2.k+1_; 1), …, (*c*_n k+1:n_, *n* − *k*). Let *P* denote the concatenated sequence of all pairs generated by the strings in *R* and let *P*_w_ denote the sequence obtained by first taking the subsequence of pairs in *P* whose first component equals *w* and subsequently projecting each of those pairs to it’s second component (thus retaining a sequence of positions where *w* appears in the reads in R). Let *H*_*w*_ denote the frequency histogram of *P*_*w*_, i.e. the function mapping each position *w* appears at in any read to the number of times it appears in that position. Given the discussion of subsection 2.1 we can assign a score to each pair of *k*-mer and position pair. It may at first seem tempting to express this as a conditional probability

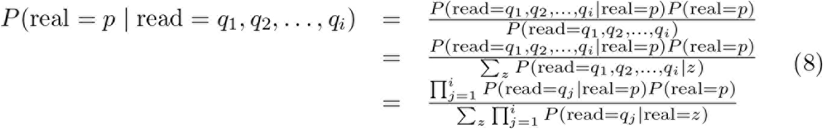

Where real = *p* denotes the event of having the real position *p* for a base and read = *q*_1_, *q*_2_, …, *q*_*i*_ the event of seeing the base at positions q_1_; q_2_,…,q_*i*_ in reads. The function *P*(read = *q*_*j*_ | real = *p*) is easily expressed using the function *P*_*t*_ defined above. This approach however has several drawbacks. Firstly we do not know the function P (real = *p*). We could obtain an approximation by assuming a constant value inside a window of reasonable position values though. Secondly and more severely this approach suffers from it’s dependence on a precondition we do not have, namely that all positions for a k-mer in the input stem from a single instance of the *k*-mer in the real sequence. This precondition does not hold for two reasons. Firstly a *k*-mer may appear multiple times in the assembled sequence. Secondly the appearance of a *k*-mer may be due to an error. Both of these lead to position sets which are incompatible with obtaining a non-zero probability out of *P*(real = *p* | read = *q*_1_; *q*_2_, …, *q*_*i*_) and we would obtain a zero probability for each real position even though a subset of the reported read positions would give evidence for a non-zero probability for certain real positions. We avoid this issue by instead of trying to obtain *P*(real = *p* | read = *q*_1_, *q*_2_, …, *q*_*i*_) computing a score for each pair of a *k*-mer and a real position it may occur at. To this end we define the function kscore by

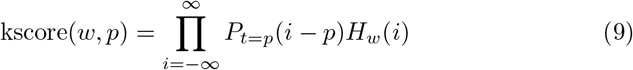

For a *k*-mer *w* and a position *p*. kscore is easily evaluated due to the finite support of both *P*_*t*_ and *H*_*w*_. It is resilient to single outlier positions produced by erroneous *k*-mers, as those do not harm the score of a real occurrence. As the support of *P*_*t*_ is narrow for small values of *t*, kscore produces separate peaks for all but very close repetitions of a *k*-mer in our application. Having separate peaks is not crucial for our application though.

### 2.6 Extracting Windows from Alignment Piles

As we want to compute consensus sequences over small windows of size 100 base pairs or less, we need to obtain the respective read data for such windows. Given an alignment pile for a read *A* it is a simple task to extract a sub pile starting and ending at any given position of *A* using the edit scripts contained in the alignment tuples. This yields the information which read data we need to use to build a De Bruijn graph. Figure 7 depicts a window on an alignment pile. For computing consensus sequences we use only read data from reads which fully span a window, i.e. not such which start or end inside the window. In practice it has proven to be beneficial to extract overlapping windows instead of partitioning a read into a sequence of non-overlapping intervals. Once we have extracted the read data for a window, we can build it’s De Bruijn graph.

**Figure 7:**
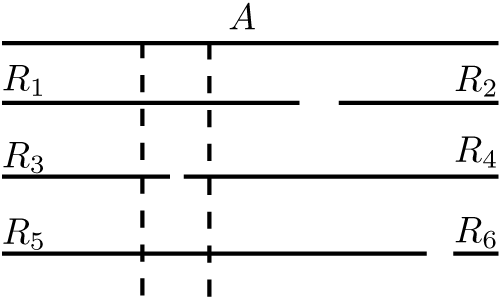
Window (dashed lines) on an alignment pile for a read *A*. Reads *R*_1_, *R*_3_, *R*_4_ and *R*_5_ overlap the window. Reads *R*_1_ and *R*_5_ span the window.

### 2.7 Estimating Consensus Length

One of the crucial tasks for finding a good reconstruction for a window is having a good estimate of the length of the reconstructed sequence. Note that due to the fact that the insertion and deletion error rates for long read sequencing are often different, the average length of the input data is not a good estimator of the consensus length. For PacBIO data, where we usually see *p*_*i*_ > *p*_*d*_, the consensus sequences are usually shorter than the average of the read data. A better method is to consider the set of last *k*-mers appearing in the reads. Let *w* be the one with the maximum frequency, i.e. there is no other k-mer w' s.t. more reads end on *w*' than on w. Let *p* = arg max_i_ kscore(*w*, *i*), i.e. the position *p* maximising the kscore function for *w*. Then *p* + *k* is a good estimate of the length of the consensus sequence. We call the maximal interval 
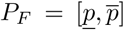
 s.t. kscore(*w*, *p*') ≠ 0 for *p*' ≠ *P*_F_ the interval of feasible lengths for a window.

### 2.8 Paths through the De Bruijn Graph

Any path through a De Bruijn graph can be specified by the sequence of nodes *v*_1_, *v*_2_, …, *v*_*n*_ it visits. We assign the weight 
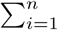
 kscore(*v*_*i*_; *i* − 1) to the path *v*_1_, *v*_2_, …, *v*_*n*_. An obvious choice for a starting node of a path is the most frequent first *k*-mer found in the reads. At low depths it may pay off to try several starting nodes though, as the true first *k*-mer may be no more frequent than any error k-mer. As the support of *P*_*t*_ widens and thus the expressiveness of the scores yielded by kscore decreases with increasing position it is beneficial to keep the positions as low as possible. For a single path there is no way to achieve this, but we can traverse the graph in two directions instead and look for points where we can join up the paths originating from the read fronts and read backs. As we already noted for estimating the consensus length, a suitable start node from the back of the reads is the most frequent last *k*-mer found in the read set. As for the case of starting from the read fronts it may be beneficial to try multiple alternatives in the case of low depth. When we join up two paths originating from the read fronts and read backs the combined path weight is the weight of the path from the front plus the weight of the path from back minus the weight assigned to the last node of the front originating path (as otherwise we take that node into account twice). Let 
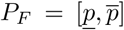
 denote the interval of feasible lengths for the window underlying the De Bruijn graph considered. We traverse the graph from the back of the reads until we reach a path length of 
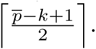
. Let Q denote the set of paths obtained in this way. Then we traverse the graph from the front of the reads up to a length of 
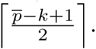
. As soon as we reach a path length of at least 
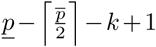
 we try to connect up the path with the paths in *Q*. Figure 8 depicts the traversal strategy. As a practical (and heuristical) optimisation we do not follow a path any further (from the front as well as from the back) if we have already found a given number (say c) of higher scoring paths of the same length. This is easily achieved by keeping a (min) heap for each length storing the up to c top scoring paths of that length found so far. Assume we have a new path of length *l*. If the heap for length *l* contains less than *c* paths so far, then we add the new path. Otherwise the heap is full. If the new path has a score not exceeding the score of the lowest scoring path in the heap, then we discard the new path. Otherwise we discard the lowest scoring path in the heap and add the new path for length *l*. In addition we do not consider all front and back path pairs possible but only a given number of highest scoring ones. To this end let *Q* = {*Q*_1_, *Q*_2_, …, *Q*_n_g. Let first(*Q*_i_) denote the first *k*-mer in the path *Q*_i_ (i.e. the last *k*-mer which was added starting from the back of the reads), and let pscore(*Q*_i_) denote the score of the path *Q*_i_ as defined above. Let *Q*_w_ denote the set of *Q*_i_ in *Q* s.t. first(*Q*_i_) = w. Consider any linearization (*Q*_w1_, *Q*_w2_, …, *Q*_w__m_) of *Q*_w_. We assign an integer rank(*Q*_w__j_) 2 f0; 1; m 1g to each *Q*_w__j_ s.t. rank(Q_w__a_) < rank(*Q*_w__b_) if pscore(*Q*_w__a_) < pscore(*Q*_w__b_) or if pscore(*Q*_w__a_) = pscore(*Q*_w__b_) and *a* < *b*. The rank function induces a strict ordering on *Q*_w_ in terms of the scores of it’s elements. Now we index the sequence rank(*Q*_w1_), rank(*Q*_w2_), …, rank(*Q*_w__m_) for range maximum queries (see e.g. [8]) and range quantile queries using a wavelet tree (see [9]). The range maximum queries allow us to find one highest scoring path in constant time. The range quantile queries facilitate finding the next lower scoring path relative to a given path in time O(log *m*). For enumerating complete paths in decreasing order of combined score consider the set of front originating path *P* = {*P*_1_, …, *P*_*n*_}s. For each *P*_*i*_ let last(*P*_*i*_) denote the last *k*-mer in *P*_*i*_ and let pscore * (*P*_*i*_) = pscore(*P*_i_) kscore(last(*P*_i_); |*P*_i_| k 1) the adapted path score for P_*i*_ (remember we subtract the score for the last *k*-mer as to not consider it twice). Now, using another heap, it is simple to enumerate the sequence of pairs (*P*_*a*_, *Q*_b_) s.t. last(*P*_*a*_) = first(*Q*_b_) in decreasing combined score order s.t. retrieving the next element takes time *O*(log max{|*P*|, |*Q*|}). If there is insufficient depth or an excessive error rate then we may encounter the case that there is no path from the chosen start node to the chosen end point. In this case we cannot produce any consensus sequence.

**Figure 8:**
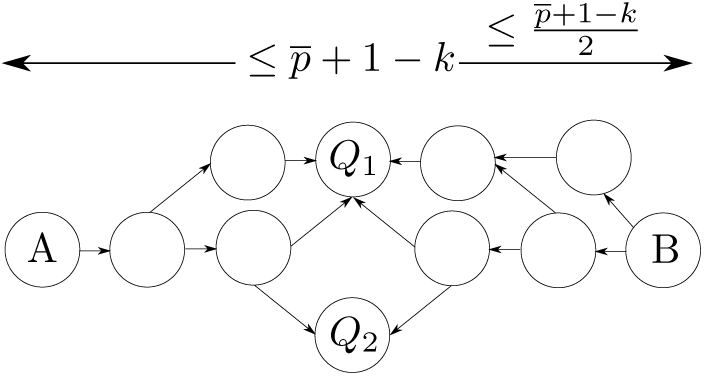
De Bruijn graph traversed from front of reads (node A) and back of reads (node B)

### 2.9 Selecting Consensus

We constructed a De Bruijn graph for a window of an alignment pile of a read A. This De Bruijn graph is based on a sequence of strings *S*_1_, *S*_2_, …, *S*_*n*_, the rst being a window from A and the latter windows from other reads aligning with A. The graph traversal will in general not yield a single path but a set of paths. Each of these paths spells out a possible consensus sequence. Let {*C*_1_, *C*_2_, …, *C*_*m*_} denote the set of possible sequences. For each *C*_i_ we compute optimal pair wise (global) alignments between *C*_*i*_ and the strings *S*_*i*_. We choose the string *C*_*i*_ yielding the lowest accumulated error as the consensus of the *S*_*i*_.

### 2.10 Combining Window Consensus to Read Consensus

Up to this point we have computed consensus sequences of small windows on a read alignment pile. We combine these windows by first aligning each window consensus to it’s respective window on the *A* read. Let the window start at base number *b* in *A* and let *S* = *s*_1_, …, *s*_n_ denote the resulting edit script for transforming the window of *A* into the consensus. We assign position pair (b + *i*; 0) to the *i*’th non insertion in *S* and position pair (*b* + *i*, − *d*) to the *d*’th insertion before the i’th non-insertion counted from right to left (i.e. if *S* starts with two insertions and a match then the first three position pairs are (*b*, −2), (*b*, −1) and (*b*, 0)). Each position pair is annotated with the respective symbol of the window consensus (or D if the edit script has a deletion for the position) thus generating pairs of position pairs and symbols. We then sort these pairs in lexicographical order by the position pair component. The final consensus is obtained by using a simple majority vote for each position pair. In case no symbol has a clear majority we arbitrarily choose the first eligible symbol in the order *A*, *C*, *G*, *T*, *D*. We then combine each sequence of consecutive position pairs without gaps into a consensus sequence, i.e. a read may have more than one stretch of consensus if the path generation fails for some intermediate windows. Majority symbols *A*, *C*, *G* and *T* are retained, positions with majority symbol *D* are dropped.

### 2.11 Patching in Missing *k*-mers

For low depth it often happens that the De Bruijn graph does not yield a path from a chosen start to a chosen end node. In this case it sometimes proves effective in practice to patch in *k*-mers which may be suspected as missing. If for instance we have *k*-mers *w* and *v* s.t. *w* = *c*_1_*c*_2_… *c*_k_ and *v* = *c*_3_*c*_4_ … *c*_k_c_k+1_*c*_k+2_ then we insert *u* = *c*_2_*c*_3_ … *c*_k+1_ at a position *p* + 1 where *p* is any position in which *w* appears in any read if *u* does not already appear in the graph.

### 2.12 Estimating Error Probabilities

Our modelling and scoring schemes rely heavily on the error probabilities p_*i*_, *p*_*d*_ and *p*_*s*_, so it is crucial to have good estimates of these probabilities. One way to obtain these values is to resequence a species with a known reference genome, map the reads to this reference and extract the error probabilities from the resulting alignments. Another way which works well in practice is the following. Consider a short window (≤ 40 base pairs) on an alignment pile of high depth d (≥ 100). We build a De Bruijn graph for the window. Then for each node we remove all edges but the one to the most frequent successor node (i.e. the one to the *k*-mer among the successors which appears most often in the reads). We then traverse the graph from the most frequent starting to the most frequent last *k*-mer in the window. If there is no such path, then the window most likely contains a repeating *k*-mer and we disregard it. Otherwise the path obtained spells out a consensus candidate. Assume there would be a repeating *k*-mer in the true sequence of the window. The probability of seeing this repeating k-mer twice in any of the reads is at least (*p*^*k*^)^2^, where *p* is the probability of a correctly reported base. This implies we have a chance of at least 1 − (1 − (p^2*k*^))^*d*^ to see the repeating *k*-mer twice in a single read inside the pile. For *p* = 0.14 as we typically find it in PacBIO data and *k* = 8 and *d* = 100 we obtain the probability 0.9996 and are thus almost sure to see this event. When we detect a repeating *k*-mer then we again disregard the underlying window. If we did not disregard the window because we consider it as containing a repeating *k*-mer, then we consider the consensus as similar to reference quality and align the *A* read fragment underlying the window to the consensus. We accumulate alignment statistics data (number of matches, mismatches, insertions and deletion) for a large set of windows. Finally we compute average error rates from the accumulated data to obtain estimates for *p*_*i*_, *p*_*d*_ and *p*_*s*_. Figure 6 shows that there is an ample number of windows in species like E. coli and D. mel for this approach to work. In both cases more than 80% of the windows of length 40 can be traversed without encountering a branching node.

### 2.13 Intrinsic Quality Values

Intrinsic quality values are a concept which has already appeared in the Dazzler suite. The quality of a read region (usually panels of 100 bp on a read) is estimated by considering the error rate at which other reads align to it in that region. More precisely given a depth parameter d the intrinsic quality value for a position interval [*it*; (*i* + 1)*t*) of a read *A* is obtained by taking the top *d*/4 (in terms of error rate) reads aligning to *A* in [*it*, (*i* + 1)*t*), summing up the number of errors between *A* and those reads in [*it*; (*i* + 1)*t*) and dividing it by *d*/4. If less than *d*/4 reads align to *A* in an interval than the read is considered as broken in this interval. As the intervals have fixed size the average number of errors of the top *d*/4 aligning reads can be transformed into an average error rate. Figure 9 depicts a graphical representation of intrinsic quality values on a read. In our context the main benefit of intrinsic quality values is to provide a hint as to whether an alignment between two reads ends although none of the two reads ends because either (at least) one of the reads has a sharp drop in quality or both reads have good quality but no longer correlate. The latter case usually signifies that two reads align due to a common repeat region, but the alignment ends because the regions the reads stem from are not correlated beyond this repeat region.

**Figure 9:**
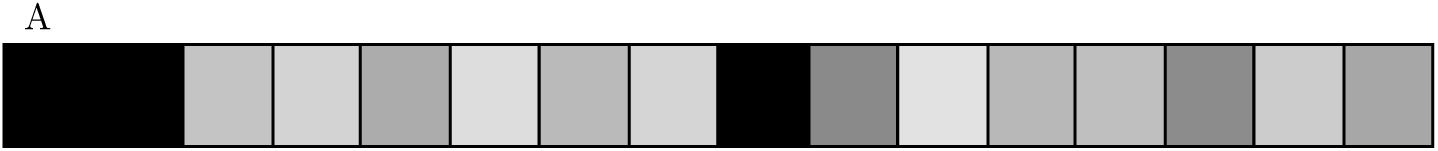
Intrinsic quality values for read *A*. Lighter colours denote better quality, darker ones worse quality.

### 2.14 Simple Repeat Detection

As mentioned in subsection 2.3 we need to distinguish between the case of simulated data giving us the opportunity to obtain perfect alignment piles and the case of real data where we start off from a set of local alignments between reads, not all of which will represent true overlaps in the genome. The non true alignments in alignment piles are induced by repeats in the underlying genome.

We will present two simple methods to remove a part of the non true alignments from alignment piles which contain all significant local alignments between read pairs. To this end assume we are given the following:

1. a set of reads *R* = {*R*_1_, *R*_2_, …, *R*_r_}
2. an alignment pile *P*_i_ for each read *R*_i_ containing all the significant local alignments between *R*_i_ and other reads in *R* (for instance compute using DALIGNER [21])
3. intrinsic quality values for each read *R*_i_.

We say that an alignment *S*_i_ between two reads *R*_*i*_ and *R*_*j*_ has a proper left (right) end, if S_*i*_ extends to the left (right) until either *R*_*i*_ or *R*_*j*_ ends or either *R*_*i*_ or *R*_*j*_ has low quality (as signified by the intrinsic quality values for *R*_*i*_ and *R*_*j*_). We call an alignment *S*_i_ between *R*_i_ and *R*_j_ proper, if it has both a proper left and right end. Figure 10 depicts a proper and an improper alignment between two reads A and B. Note that although the alignment *O*_1_ in Figure 10 is a proper alignment according to this definition, the fact that the two reads also share an improper alignment O_2_ gives evidence that *O*_1_ should also be disregarded. If a read *A* shares an improper alignment *S* with a read *B* s.t. *S* start at base *b* and ends at base *e* in *A*, then we say that *S* gives evidence for a repeat [*b*, *e*] on *A*. *A* position *p* on a read *A* is inside a *c*-repeat, if at least *c* alignments between *A* and other reads give evidence that *p* is in a repeat region. We could consider 1-repeats as sufficient to declare regions in a read as repetitive, but as we are handling noisy data it is safer to use a somewhat higher threshold *c* in practice. As a first method to remove non true overlaps, we remove all such alignments between a read *A* and a read *B* s.t. *A* and *B* share at least one improper alignment. Note that for reads without low quality regions this would essentially boil down to removing all alignments which are not of the suffix/prefix type, i.e. at both ends of the alignment a suffix of one read aligns to the prefix of another read. As however we often find low quality regions in real data, the more general approach proves to be useful in practice.

**Figure 10:**
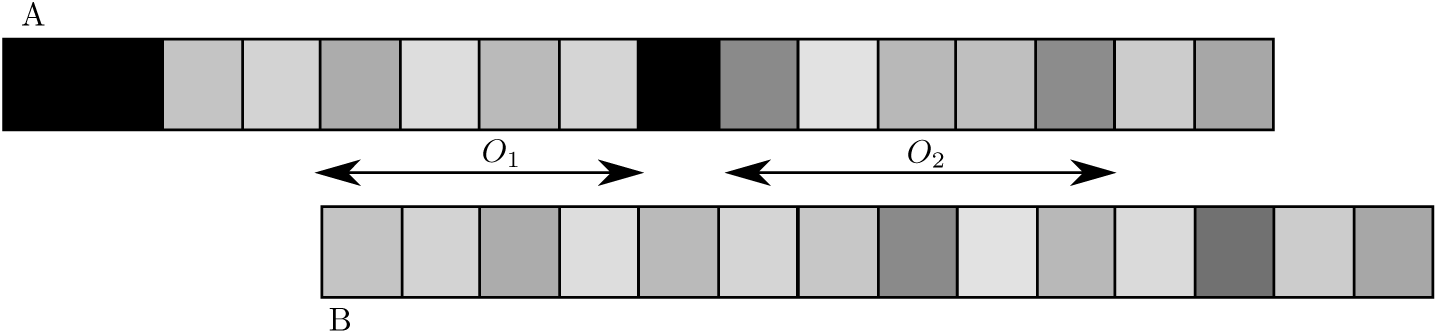
Proper and improper alignments: the reads *A* and *B* share the local alignments *O*_1_ and *O*_2_. *O*_1_ is a proper alignment. It has a proper left end because it reaches the left end of *B* and a proper right end because it reaches a low quality region of *A* on the right. *O*_2_ is an improper alignment. Whilst it has a proper left end as it extends to the edge of a bad region in *A*, it ends on the right without extending to a read end or bad quality region. This suggests *A* and *B* are no longer correlated although they have good quality.

The approach so far only removes improper alignments. There are however non true repeat induced alignments which do not produce improper alignments. One sub class of these can be detected, as we describe in the following. Let A and C be two reads featuring a suffix/prefix alignment where a prefix of *C* aligns to a suffix of *A*. The alignment of *A* and *C* is covered by a repeat *R* in the underlying genome. *A* has a segment called *U* to the left of *R*. *C* has a segment called *V* to the right of *R*. Figure 11 depicts this scenario. The picture has *R*', which may be a prefix of *R*. Note that the alignment between *A* and *C* is proper if *A* and *C* both are of good quality. It ends on the left because it extends to the left end of *C* and on the right because it extends to the right end of *A*. Assuming the left context *U* of *R* in *A* and the right context *V* of *R* in *C* are sufficiently long we can check whether *U* and *V* are compatible contexts for *R* or, in other words, whether the combination *URV* appears in the underlying genome. For detecting this scenario we first need to mark *c*-repeats for some suitable *c* in the reads using intrinsic quality values. Then, given the marked repeats, we check for a given read *A*, whether it’s suffix (the prefix case is symmetric) aligns to another read *B* which has a repeat region *R* covering a suffix of *A* (see Figure 11). If this is the case, then A ends in a repeat. Now we check for reads *C* which have a prefix aligning to a suffix of A s.t. the alignment is covered by the marked repeat. W.l.o.g. let *C* overlap a suffix of *A* as depicted in Figure 11. We check whether there are reads *D*, which properly align to a suffix of A, extends at least a given number of bases into *U* and has an overhang of a given number of bases to the right of A. If there are any such reads *D*, but *C* does not properly align to any of them because the right context of *R* found in *C* is not compatible with any of the right contexts found in the *D* reads, then we discard the alignments between *A* and *C*.

**Figure 11:**
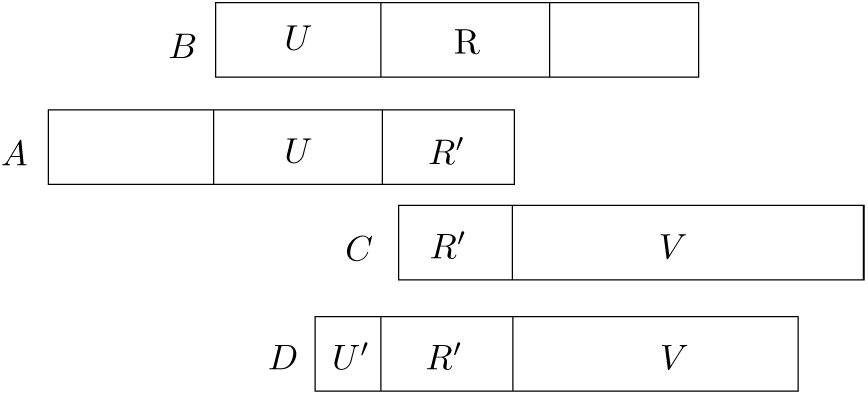
Read *B* contains a marked repeat *R*. A suffix of *A* aligns to a prefix of *B* and the alignment is covered by the repeat *R*. A prefix of another read *C* aligns to a suffix of *A* and this alignment is covered by the repeat *R*.

The two filtration steps described above remove a large number of spurious alignments not stemming from real overlaps. Some however, particularly those due to reads which are completely contained in repeat regions, remain. Removing these requires a deeper analysis of the read data beyond this algorithmic work on read consensus computation. One feasible way is to perform repeat separation by computing a preliminary consensus sequence and then separating reads in groups which disagree with this consensus in a consistent way. We will address this open problem in upcoming work.

## 3 Experimental Results

We have implemented the approach described above in a program called daccord. It is freely available at https://github.com/gt1/daccord. For testing we used simulated as well as real long read data. Simulated reads were produced using the PacBIO read simulator PBsim (cf. [22]) with mean length of 15000 bp and mean accuracy of 85%. For simulations we used the genomes E. coli (strain K-12 MG1655, GenBank U00096.2) and Saccharomyces cerevisiae (strain S288C, NCBI accession NC 001133-NC 001148 and NC 001224, R64-2-1 20150113). We used real PacBIO data for E. coli (strain K12, data from [1]) and Saccharomyces cerevisiae (strain W303, data from [2]) as well as Oxford Nanopore data data for E. coli (strain K12 MG1655, data from [16]). All tests were performed on machines equipped with two Intel Xeon E5-2680 v3 CPUs (12 cores per CPU, hyper threading disabled) and 256GB of RAM. Tests were generally run with 24 threads. The machines were running CentOS Linux version 7.2. We first discuss the effects of some algorithmic parameters and alignment filtering on the consensus error rate achieved before comparing our approach to other published work.

### 3.1 Algorithmic Parameters

Our implementation of the algorithm described above has some user settable parameters. These concern the *k*-mer size used and window parameters. Figure 12 shows the effect of the *k*-mer size chosen on the residual average read error rate on perfect alignment piles of simulated E. coli data. Consensus for the *k*-mer intervals [7, 9] and [6, 10] is implemented by way of trying each *k* in the interval for a window and then choosing the computed consensus sequence yielding the minimum accumulated error between the reads and the consensus. We compute the residual error for a single read by first finding the longest common substring between the raw read and the computed consensus and subsequently extending this seed by running a slightly modified version of the O(*ND*) alignment algorithm (cf. [20]). The graphs show that different single values of *k* work best for different ranges of sequencing depth. A *k* value of 6 works best among those shown for a depth below 15. Finding a conserved 6-mer is more likely than a conserved *k*-mer for *k* > 6 and finding a sufficient number of conserved mers is crucial for our approach. As the depth grows finding conserved *k*-mers for higher values of *k* becomes more likely. As longer *k*-mers make De Bruijn graph traversal less ambiguous, we are more likely to find a good path. Figure 13 shows how the run-time changes with the *k*-mer size. All the curves have a local maximum for a depth below 20. This is caused by our implementation trying to reconstruct and correct as much data as possible, which requires patching in missing *k*-mers and trying several window start and end *k*-mers at low depth. As the depth increases so does the speed for higher values of *k*.

**Figure 12:**
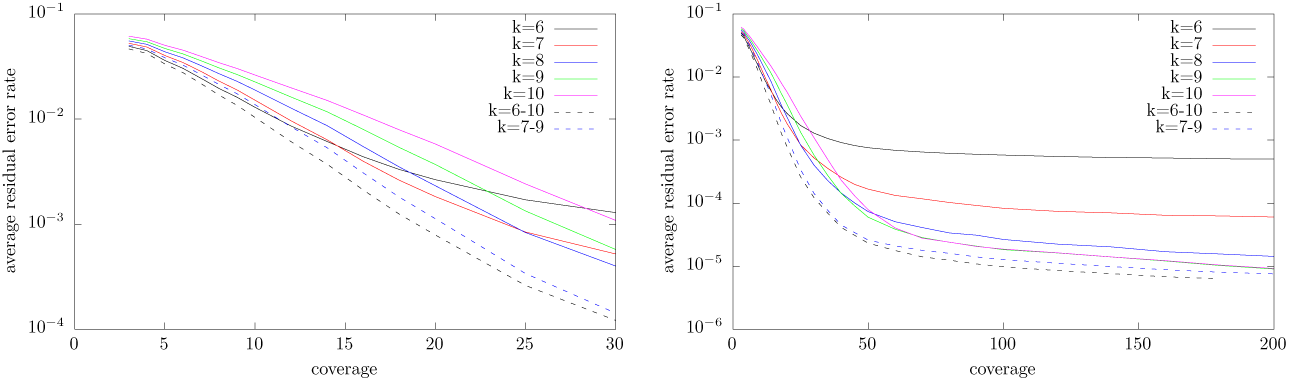
Dependence of reconstruction quality (average error rate per read) on *k* for *k* = 6; 7; 8; 9; 10 and *k* intervals [6, 10] and [7, 9] for simulated E. coli data. The left graph shows sequencing depth 3 to 30, the right graph 3 to 200. Note that the vertical axes are logarithmic.

**Figure 13:**
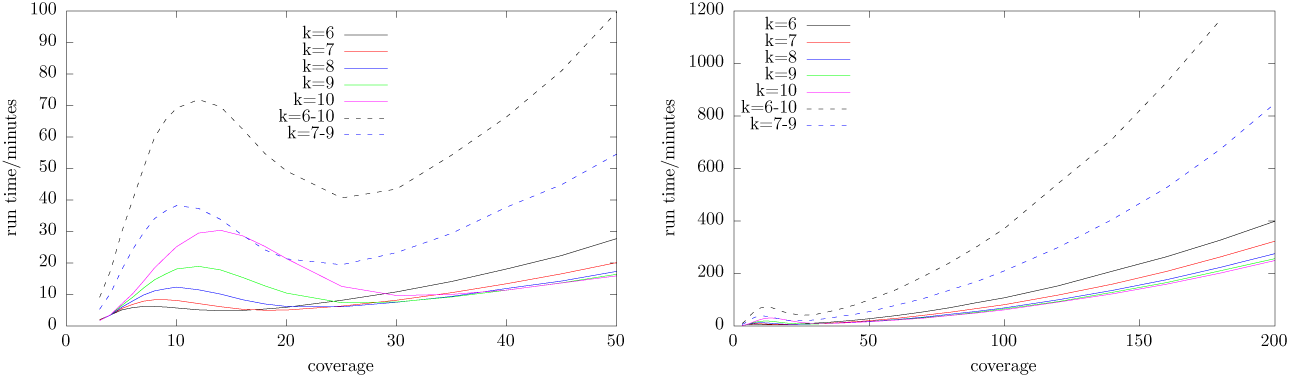
Dependence of run-time on *k* for *k* = 6; 7; 8; 9; 10 and k intervals [6, 10] and [7, 9] for simulated E. coli data. The left graph shows sequencing depth 3 to 50, the right graph 3 to 200.

The situation is similar for yeast as shown in Figure 28 (accuracy) and Figure 29 (run-time). We do however see a more pronounced benefit *t* for larger *k*-mer sizes in the accuracy graph. A *k*-mer size of 10 beats the smaller *k*-mer sizes for a depth as low as 50.

Figure 14 shows the effect of the window parameters chosen on reconstruction quality for *k* = 8. For parameters w and a the window coordinates used are [*ia*; *ia* + *w*), i.e. the i’th window starts at position *ia* and has length *w*. As the graph shows it is best to use fairly small windows of size 40 for a large range of sequencing depth settings. Even smaller windows of size 20 work slightly better for low coverage scenarios, but overall 40 is a good default value. Smaller values for a improve the consensus accuracy to some degree, but this comes at the price of increased run time as more windows need to be handled. Again the situation for yeast is similar as shown in Figure 30. The main difference is that a small window size of 20 works as well as larger window sizes even for high depth.

**Figure 14:**
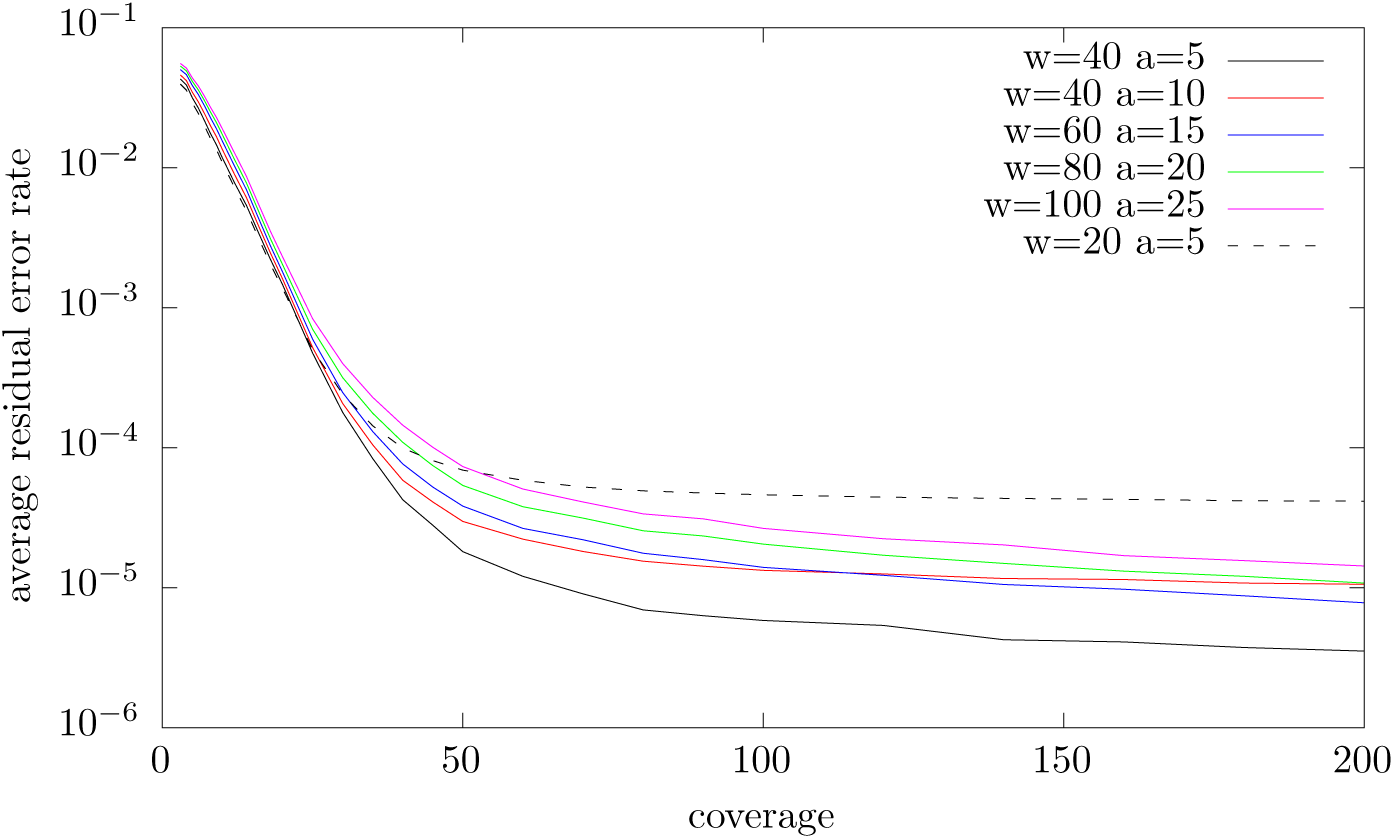
Dependence of reconstruction quality (average error rate per read) on window parameters *w* and *a*.

For the rest of the paper we fix the parameters of our runs to *k* = 8, *w* = 40 and *a* = 10.

### 3.2 Effects of Alignment Filtering

Figure 15 shows the effects of alignment filtering on correction performance. The graph compares the error rates obtained on simulated E. coli data for various sequencing depth values when using unfiltered (all significant local alignments as computed by DALIGNER are used), filtered (we use the two filters described above) and perfect alignment piles are used. The filtering shows little effect for low depth, in fact it may be detrimental. Filtering requires intrinsic qualities values and we require a depth of at least 20 to compute these. A window with insufficient depth is marked as bad, so we do not mark repeats in regions of low coverage. Filtering may be detrimental as for low depth having an alignment to a read from a highly likely repeat region is better for correction than having too little data. The effect of filtering starts to show for depth 25 and above. For depth 200 the accuracy obtained via the filtered data is an order of magnitude better (in terms of remaining errors) than the one obtained from the unfiltered data. For high depth the quality obtained for the filtered data is better than the one obtained for the perfect piles. This at first appears to be an artifact. At a closer look this can be explained by the fact that the alignments in the perfect piles continue all the way through bad read regions while DALIGNER stops an alignment if the error rate gets too high. Figure 31 shows the same type of plot for S. cerevisiae. Again filtering significantly improves the reconstruction quality. Note however that in this case the quality yielded by the filtered alignment is worse than the one for the perfect piles. This can be explained by a higher abundance of long repeats in the genome, false alignments caused by which are not removed by our two simple filters.

**Figure 15:**
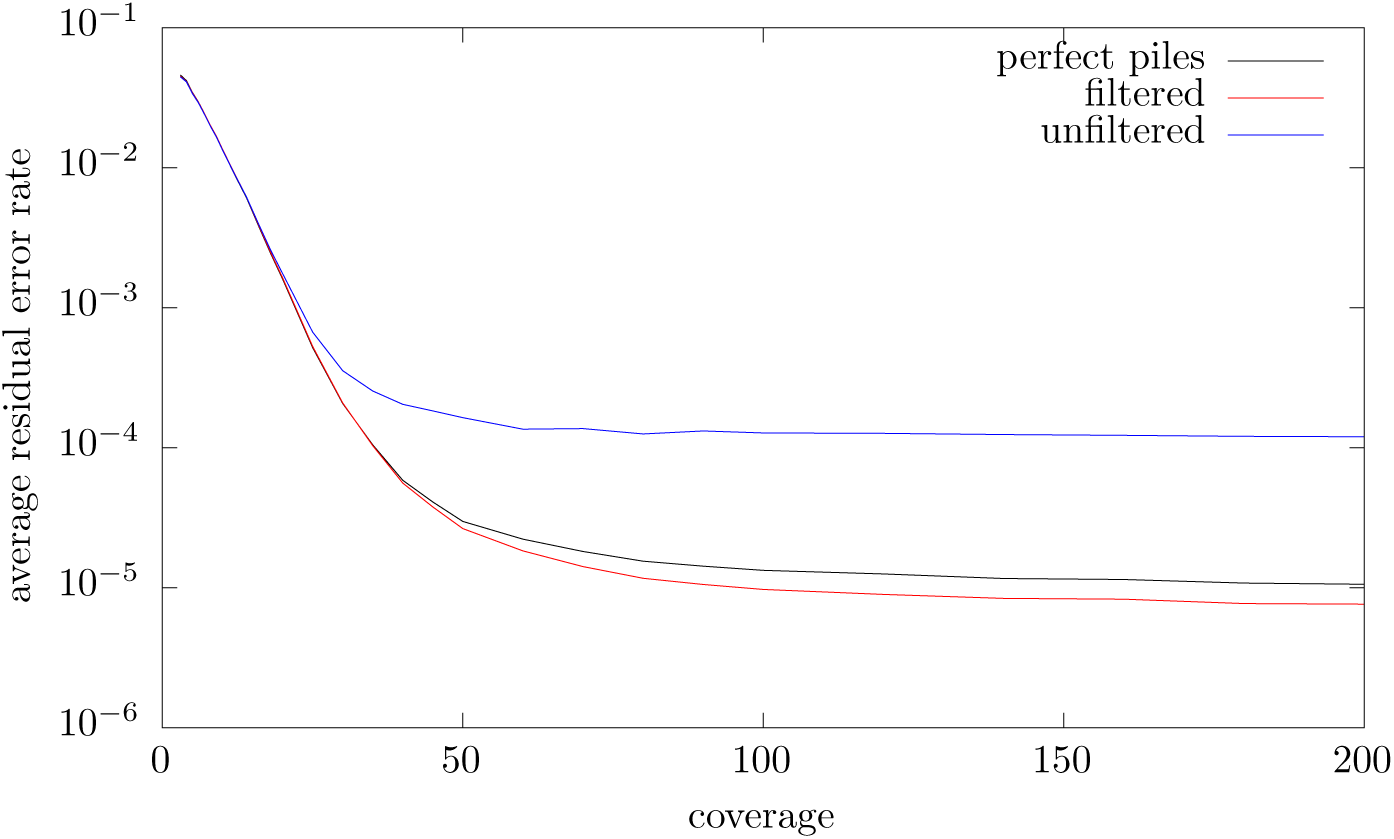
Dependence of reconstruction quality (average error rate per read) on ltering.

### 3.3 Comparison of Accuracy on Simulated Data

We first compare the accuracy of our method with two tools which work on very similar input to ours. These are namely pbdagcon (see [6]) and falcon sense (cf. [7]). pbdagcon was obtained via

https://github.com/PacificBiosciences/pbdagcon (commit f19aed1668d6ace0ab3ab4eb0e1e7d81139492a8).

We ran the program dazcon with options - c 2 - l 100 - m 10000 - j 24. This sets the minimum coverage for attempting to build a consensus to 2, the minimum length for correction to 100 and the maximum coverage used to 10000. Those parameters were chosen to make the program reconstruct/correct as much read data as possible. falcon sense was installed using FALCON-integrate available at https://github.com/PacificBiosciences/FALCON-integrate (commit 02867d30753e0510f9fed47abfb1c15fc0249f2d).

We called the program fc consensus with options −−output_multi −−n_core 24. The −−output_multi makes the program output more than one fragment of corrected data per read and again was set with the goal of correcting more read bases. Like our implementation pbdagcon and falcon sense are prepared to work with DALIGNER’s LAS file format as input. For this reason we can also observe how these tools perform on perfect alignment piles and repeat filtered alignments as described above. Figure 16 shows a comparison of daccord with pbdagcon and falcon sense for simulated E. coli data (see Figure 32 for S. cerevisiae). These graphs at first suggests a superior correction performance of falcon sense at low depth (below 16 for both). At a closer look this is however not the case. Figure 17 shows which fraction of the total bases in the read set each of the tools yields as corrected data (see Figure 33 for S. cerevisiae). At depth 10 falcon sense only outputs corrected data for half of the input data, while daccord and pbdagcon produce output for virtually all of the input data. When we compare the output of daccord with the output of falcon sense for those regions were falcon sense produces any data, then we find that for these regions our error rate is lower than the one observed for falcon sense. As this makes comparison hard, we change our comparison strategy in the following way. Both daccord and falcon sense have options to output complete sequences where uncorrected bases are given as lower case and correct bases as upper case letters. In falcon sense this can be switched on using the option −−output full. For pbdagcon (and other tools we compare to further down in the text) we map the corrected fragments onto the corresponding uncorrected reads to obtain the same structure. Furthermore for all tools we inject all input reads for which a tool has produced no output at all in the output as is. A comparison of the resulting data is shown in Figure 18 for E. coli and Figure 34 for S. cerevisiae. In this analysis daccord outperforms pbdagcon and falcon sense at every depth. A run-time comparison of daccord with pbdagcon and falcon sense is shown in Figure 19 for E. coli and 35 for S. cerevisiae. daccord is slower for low depth as it spends a lot more time trying to get a good correction based on little data. For higher depth the run-time is comparable with falcon_sense and faster than pbdagcon.

**Figure 16:**
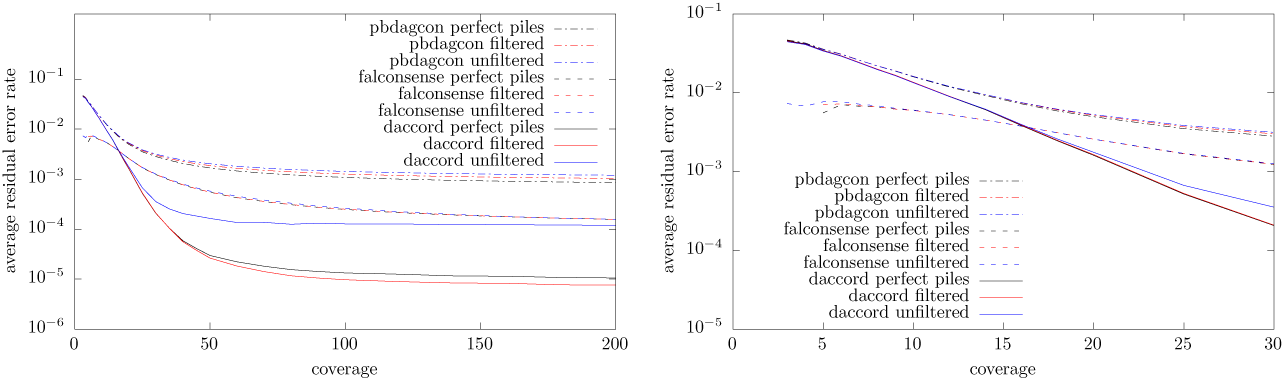
Comparison of accuracy with pbdagcon and falcon_sense

**Figure 17:**
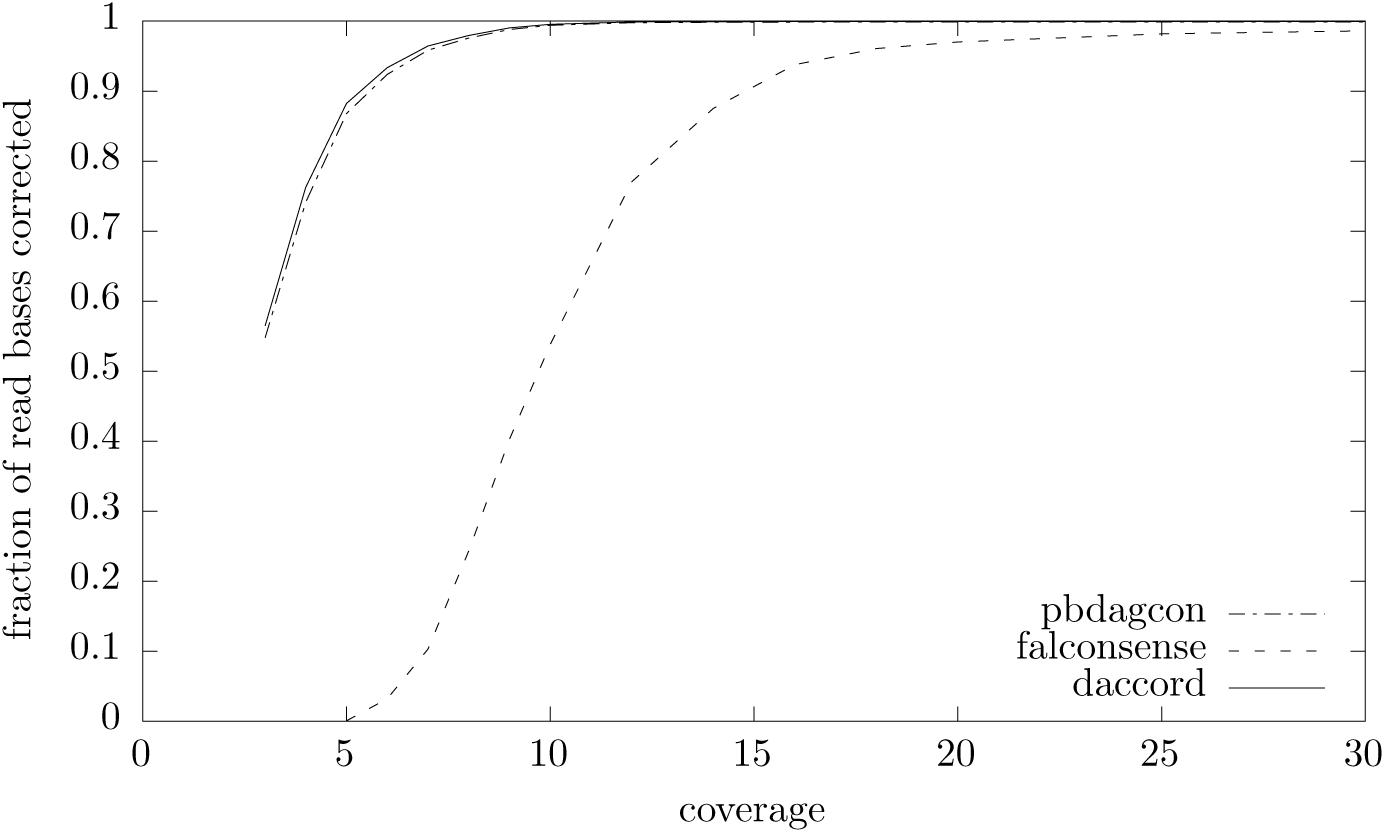
Fraction of read bases corrected by daccord, pbdagcon and falcon_sense on perfect piles of simulated E. coli data.

**Figure 18:**
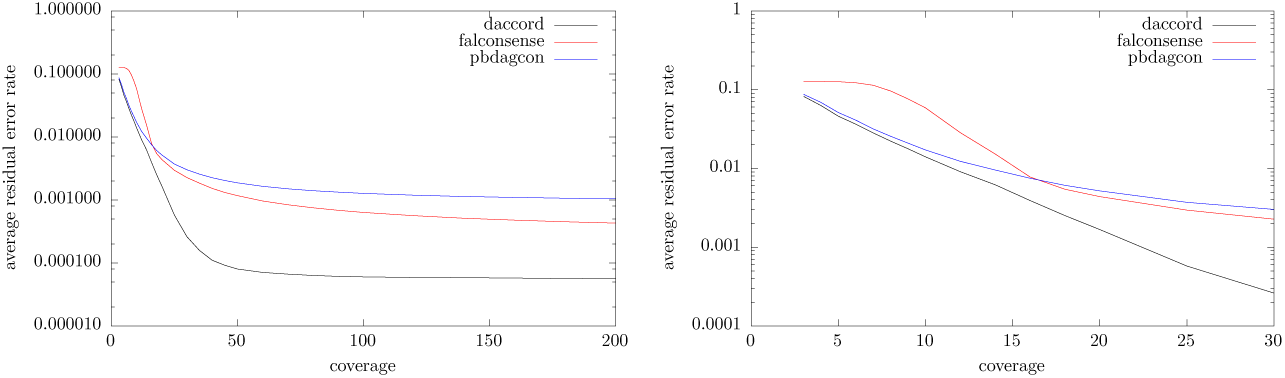
Comparison of reconstruction quality on perfect alignment piles of simulated E. coli data. Output data was inserted into the original reads for comparison.

**Figure 19:**
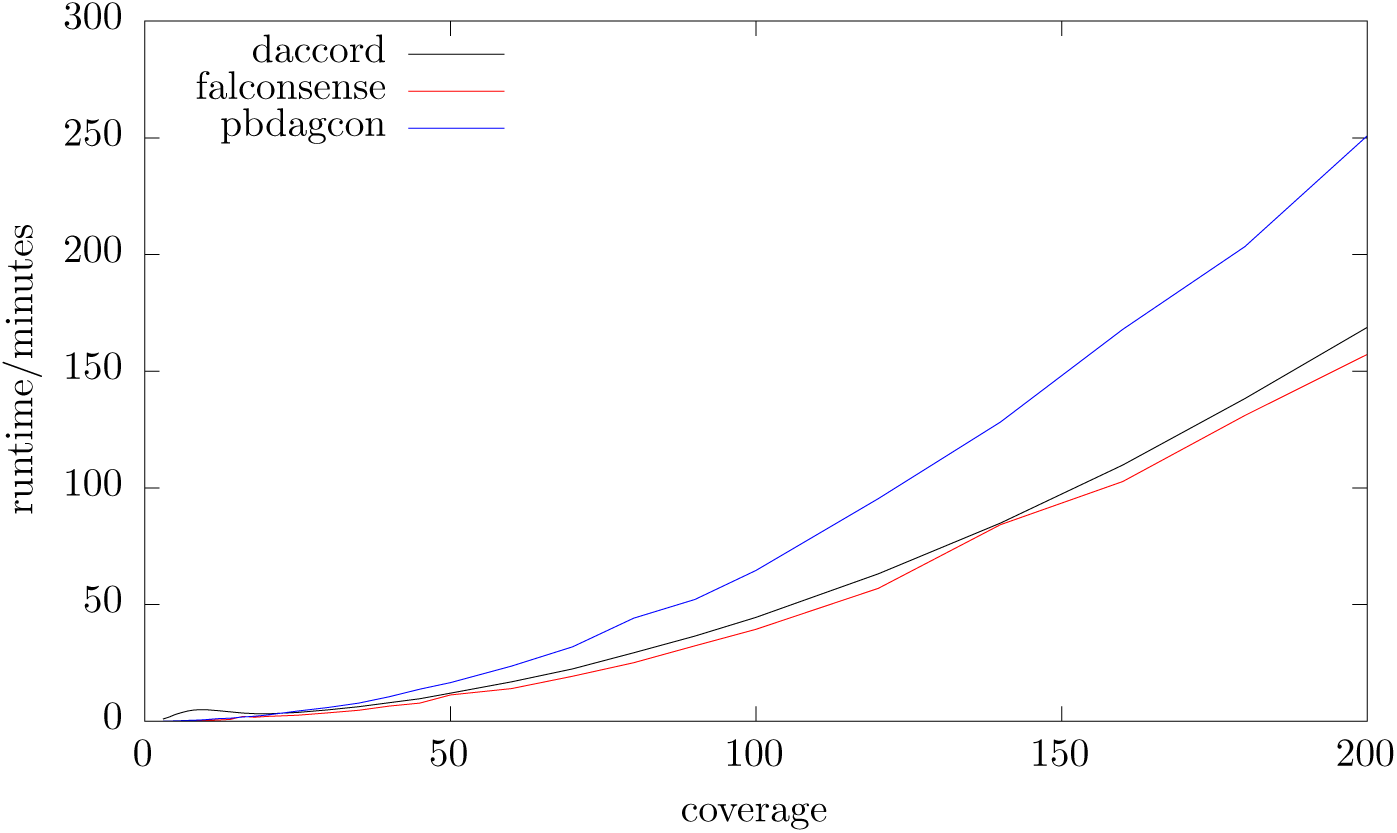
Run-time comparison with pbdagcon and falcon_sense on perfect piles of simulated E. coli data

LoRMA (see [25]) and Canu (cf. Canu [13]) are two other programs allowing non hybrid long read error correction. PBcR (see [12]) is another one, but we consider it as superseded by Canu. We used LoRMA version 0.4 and Canu version 1.4. Canu internally uses an adapted version of falcon sense. We ran LoRMA with parameters −threads 24 and Canu with −correct corOutCove-rage=300 useGrid=false −pacbio-raw and the genomeSize parameter set to the exact length of the reference in bases. We set the corOutCoverage parameter to 300 in the hope that it would correct all the reads in the input. This however did not have the desired effect. Figure 20 shows the correction performance of all the tools mentioned above (but PBcR) on simulated E. coli data (see Figure 36 for S. cerevisiae). We ran daccord, pbdagcon and falcon sense on filtered DALIGNER alignments. LoRMA and Canu were run on the corresponding raw reads. The line called *canu unfilled* in the graph does not contain data for reads Canu does not produce output for, i.e. we did not insert the raw uncorrected reads to compute the resulting error rate. We provide this line because for an assembly pipeline it may be sensible to not output a corrected version for every read as most of the corrected reads produced will be redundant, i.e. the data contained is found in other reads as well. The plot however suggests that even for a high value of 300 for corOutCoverage Canu does not produce corrected versions for most of the reads in the input at higher input depth. LoRMA requires relatively high coverage to even start any correction. The plot suggests that below a depth of 50 essentially no correction takes place. The plots in Figure 21 (E. coli) and Figure 37 (S. cerevisiae) provide more detail on this. Another problem in the output of Canu for higher depth and LoRMA can be seen in Figure 22 (E. coli) and 38 (S. cerevisiae). While for daccord, falcon sense and pbdagcon the average reconstructed fragment length quickly approaches the average length of the corresponding corrected reads, this is not the case for LoRMA and in parts for Canu. At depth 50 for instance the average length of a corrected read part produced by LoRMA is in the order of a short read length (below 300 bp). We do not provide a run-time comparison between daccord and LoRMA or Canu for simulated reads because both of the tools start from raw reads while daccord starts from pre computed local alignments on the reads. We will however be giving timings for complete correction pipelines below in the section on real read data.

**Figure 20:**
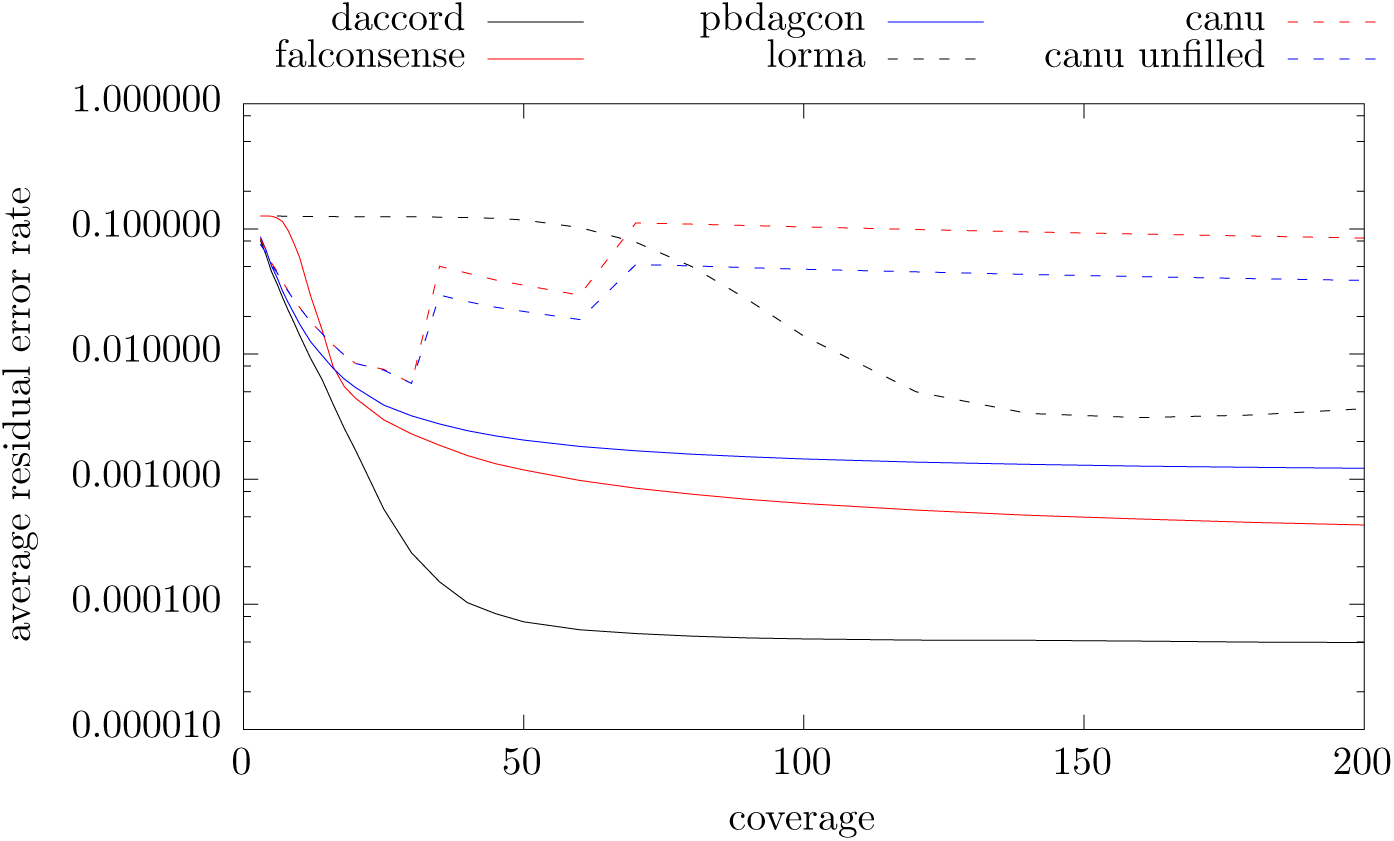
Comparison of correction performance with pbdagcon, falcon_sense, LoRMA and Canu for simulated E. coli data

**Figure 21:**
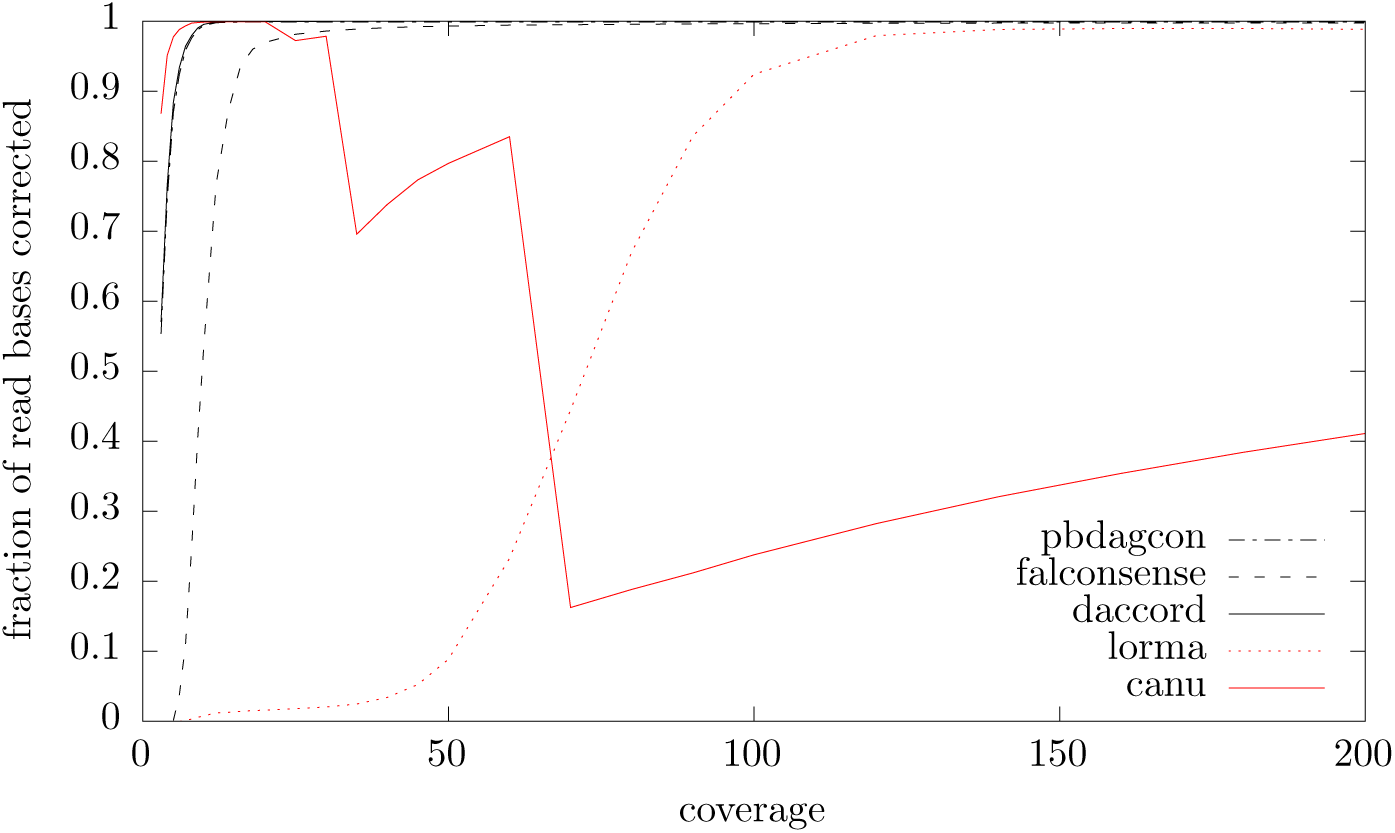
Fraction of read bases corrected by daccord, pbdagcon, falcon_sense, LoRMA and Canu on simulated E. coli data

**Figure 22:**
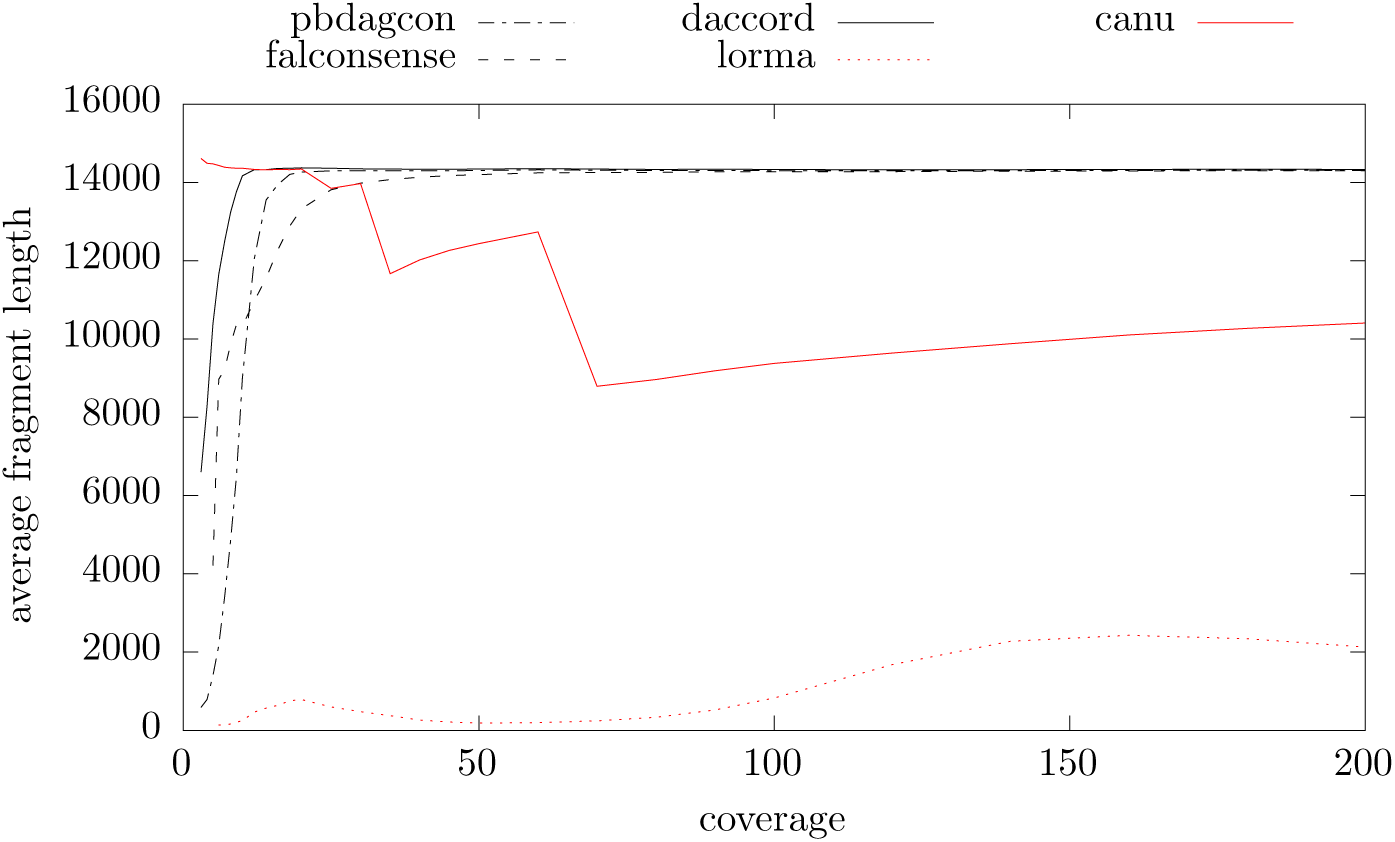
Average corrected fragment length produced by daccord, pbdagcon, falcon sense, LoRMA and Canu on simulated E. coli data

### 3.4 Comparison of Accuracy on Real Data

For real data we cannot be sure about the accuracy of a read correction process as it is often unclear where on a reference a read stems from because of repeats. In lieu of using ground truth data we can only assume a read stems from the position in a reference where it maps with a minimal number of errors. As real read data is subject to various influences like the presence of adapters, chimeric joins or stretches of (close to) pure noise, fragments of reads may not map at all and need to be clipped. For our tests we first ran the read correctors (daccord, pbdagcon, falcon sense, LoRMA and Canu), inserted the corrected sequence fragments into the raw reads, added all reads not contained in the Output of a corrector in their pure uncorrected form and finally mapped the resulting reads to the respective reference using the long read aligner damapper bwt (cf. [19, 27]). Regions clipped during mapping were disregarded as unusable. The residual error rate for a corrected read was computed based on the edit script between the mapped parts of the corrected read and the corresponding reference regions. As for simulated data the value depicted in the graphs is the average error rate for all reads.

Figure 23 shows a comparison of residual errors for real PacBIO sequenced E. coli (strain K12 MG1655, chemistry C4, enzyme P6) data. The coverage denoted on the horizontal axis is the number of sequenced bases divided by the number of bases in the reference sequence. As for simulated data daccord yields lower error rates than the other programs. Notably though pbdagcon performs better than falcon sense in the real sequencing data scenario. Table 1 shows the run-time and memory usage of the benchmarked tools on two partial data sets obtained by sub sampling. The run-time contains all steps required to obtain the corrected output when starting from raw sequence data in FastA format. In particular it contains the run-time of DALIGNER and the sub-sequent filtering steps for daccord, pbdagcon and falcon sense. Run-time and memory usage were measured using the Linux time command. Due to the way multi processing is implemented in falcon sense the time program is unable to accurately measure it’s memory usage (it only measures the memory usage of the control process but not the worker tasks performing the actual computation), so we do not provide a memory usage for falcon sense. daccord is slower on the real data than the tests on simulated data would let us expect. Figure 24 gives us a hint why this is happening. It depicts histograms showing how often reference bases were covered by read bases. The coverage is not as even as for our simulated data. For many reference bases the coverage is suffciently low to make daccord use increased effort to produce a good consensus.

**Figure 23:**
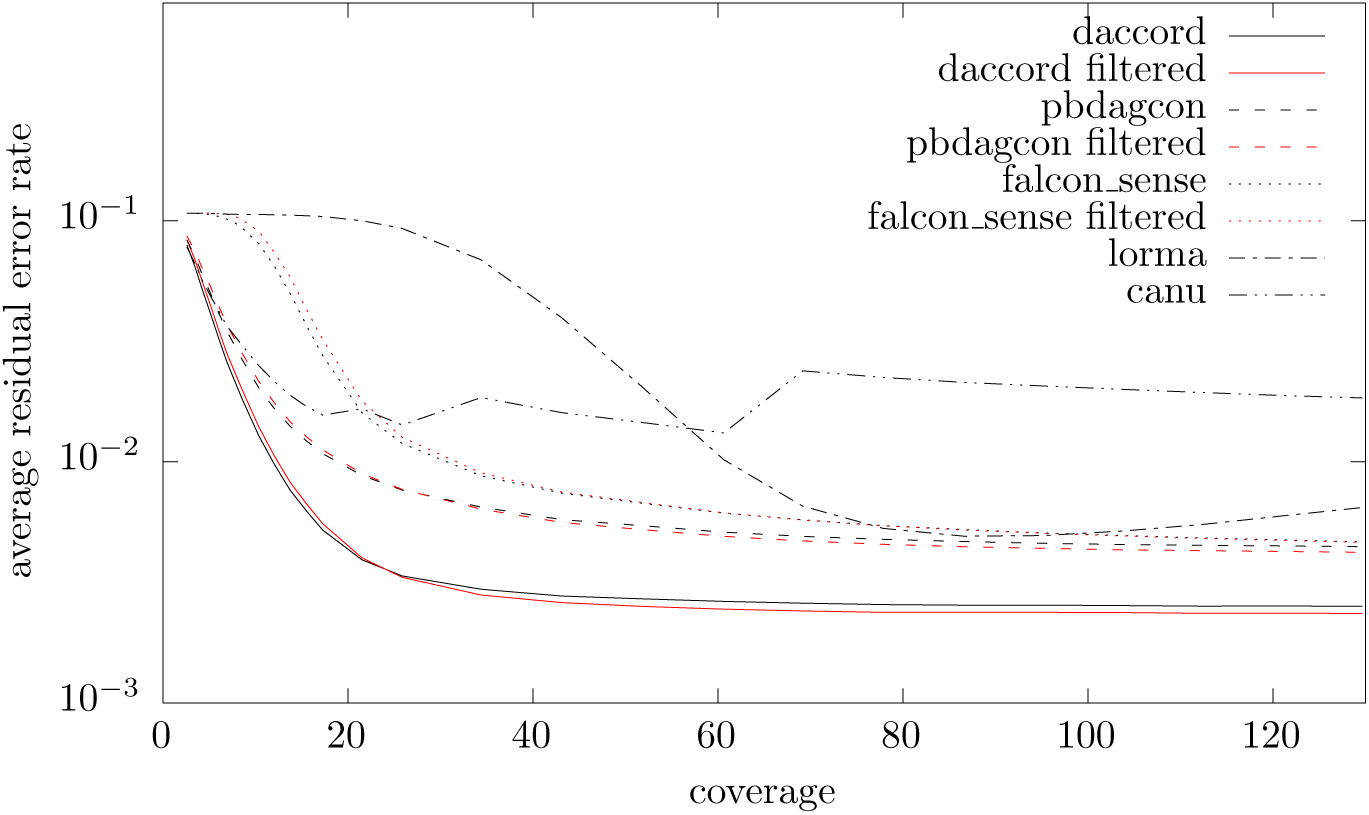
Comparison of reconstruction quality for real PacBIO E. coli data

**Figure 24:**
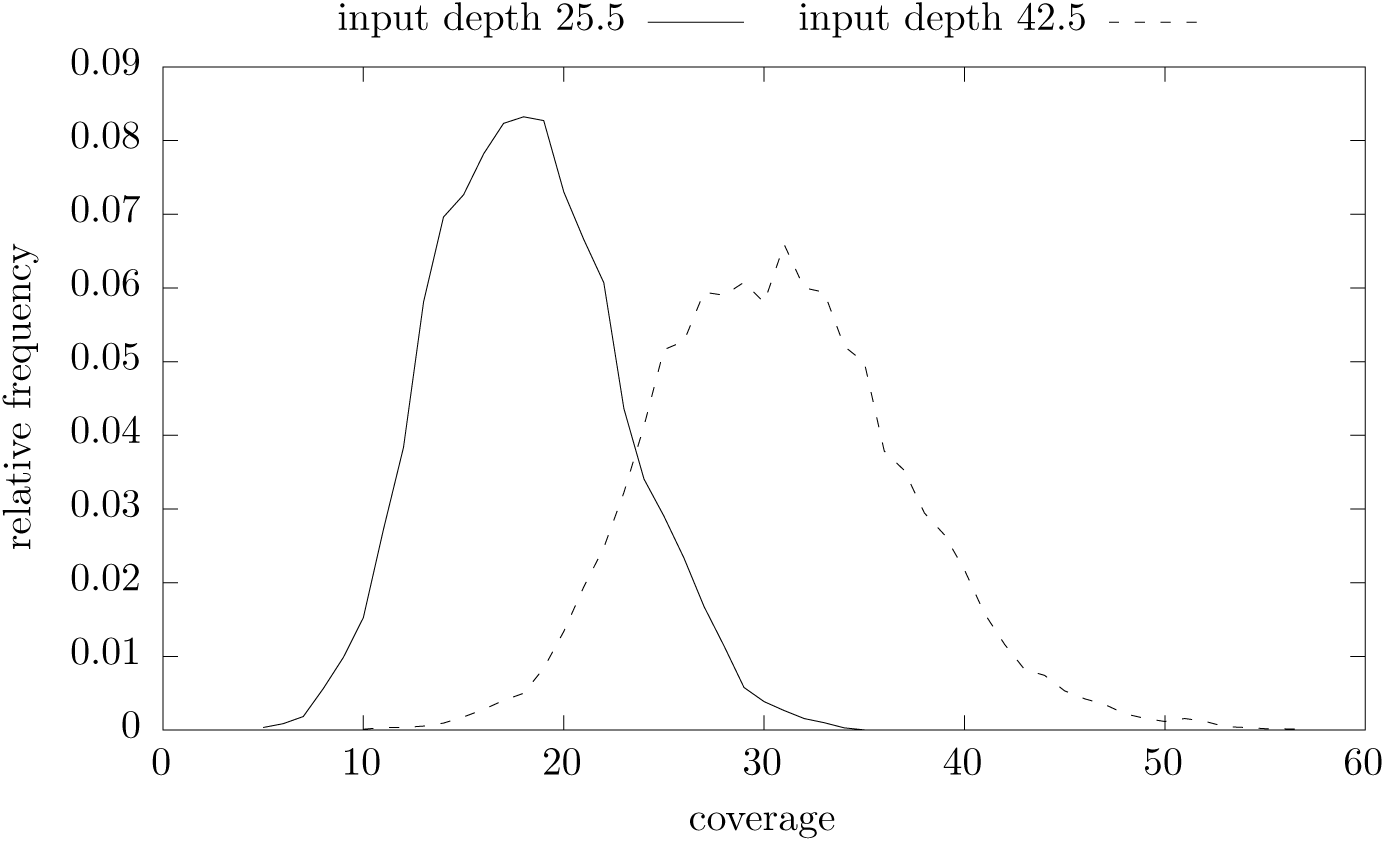
Sequencing depth histograms for real PacBIO sequenced E. coli data

**Table 1:** Run-time and memory usage of tools on real PacBIO sequenced E. coli data

|  |  |  |  |  |  |
| --- | --- | --- | --- | --- | --- |
| d=25.5 | daccord | falcon_sense | pbdagcon | LoRMA | canu |
| time | 7:13 | 3:16 | 5:37 | 4:00 | 13:29 |
| mem/GB | 5.7 | n/a | 5.7 | 55.2 | 3.8 |
| d=42.5 | daccord | falcon_sense | pbdagcon | LoRMA | canu |
| time | 13:48 | 7:31 | 14:15 | 13:22 | 17:50 |
| mem/GB | 11.8 | n/a | 11.8 | 55.5 | 4.6 |

Figure 25 shows a comparison of residual errors on yeast sequenced on PacBIO (strain W303, chemistry C2, enzyme P4). Table 2 gives run-time and memory usage for two data sets obtained via sub sampling. Canu and LoRMA are faster than the DALIGNER based pipelines on this data set. For Canu this is mainly do to the faster overlap computation employed (MHAP, see [5]). LoRMA avoids computing all pairwise overlaps, which saves time but seems to sacrifice continuity.

**Figure 25:**
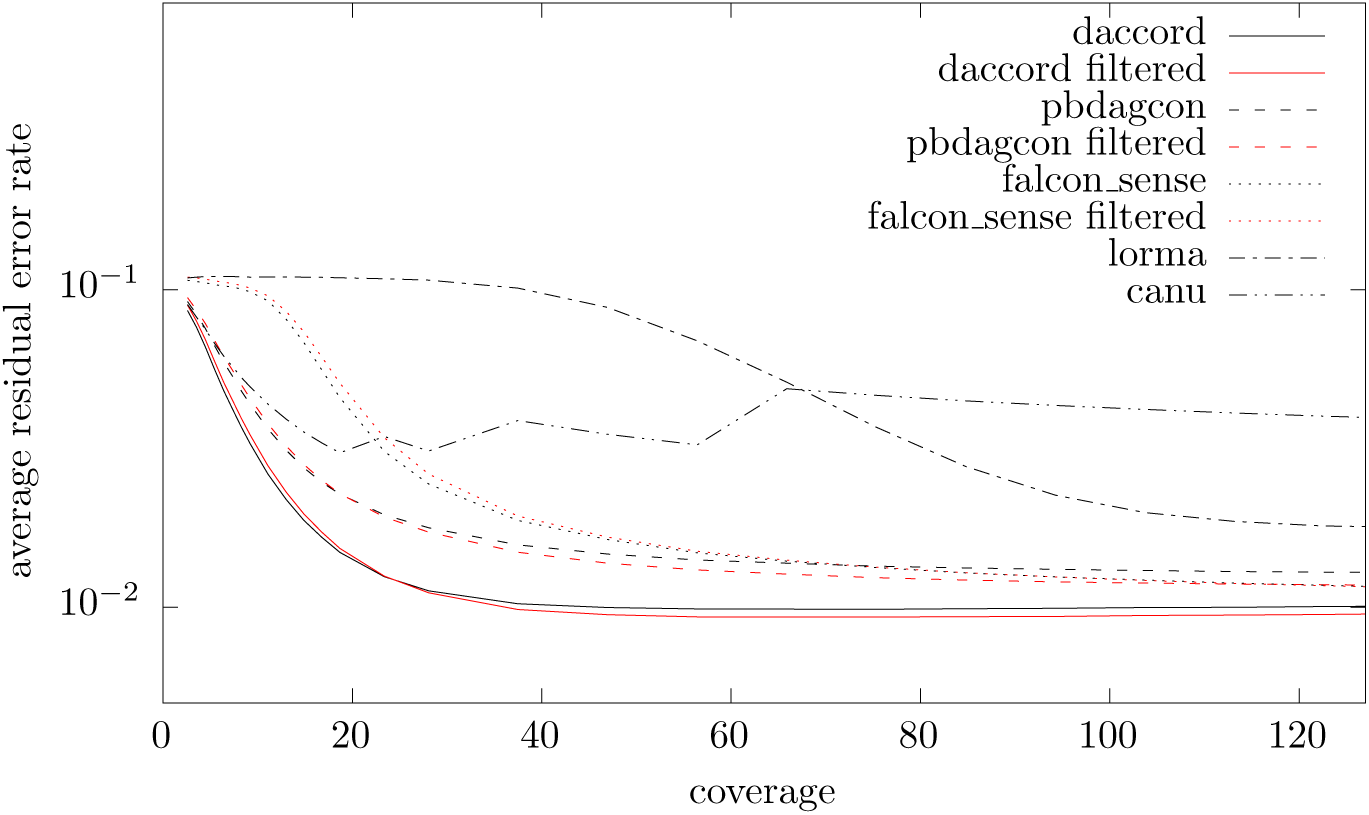
Comparison of reconstruction quality for real PacBIO S. cerevisiae data

**Table 2.** Run-time and memory usage of tools on real PacBIO sequenced S. cere-visiae data

|  |  |  |  |  |  |
| --- | --- | --- | --- | --- | --- |
| d=28.1 | daccord | falcon_sense | pbdagcon | LoRMA | canu |
| time | 58:48 | 37:39 | 1:38:01 | 33:32 | 18:27 |
| mem/GB | 20.5 | n/a | 22.0 | 55.8 | 3.8 |
| d=46.9 | daccord | falcon_sense | pbdagcon | LoRMA | canu |
| time | 2:05:14 | 1:33:29 | 4:20:08 | 1:05:05 | 30:12 |
| mem/GB | 24.8 | n/a | 39.2 | 56.6 | 4.8 |

For nanopore correction we replaced the parameter −pacbio-raw by −nano-pore-raw when running Canu. Figure 26 shows an accuracy comparison for E. coli sequenced on Oxford Nanopore (strain K12, MG1655, chemistry R9). In this plot only the higher accuracy 2D reads were used for consensus computation. daccord is also the most accurate program in this category. Table 3 shows a run-time and memory comparison for two subsets of this data set. Figure 27 and Table 4 show the same for more inaccurate 1D Oxford Nanopore data sequenced from the same E. coli strain. For this data set even at very high depth we have a residual error rate of more than 2%. This suggests effectively correcting this type of data requires a more machine specific method than our iid random event based approach.

**Figure 26:**
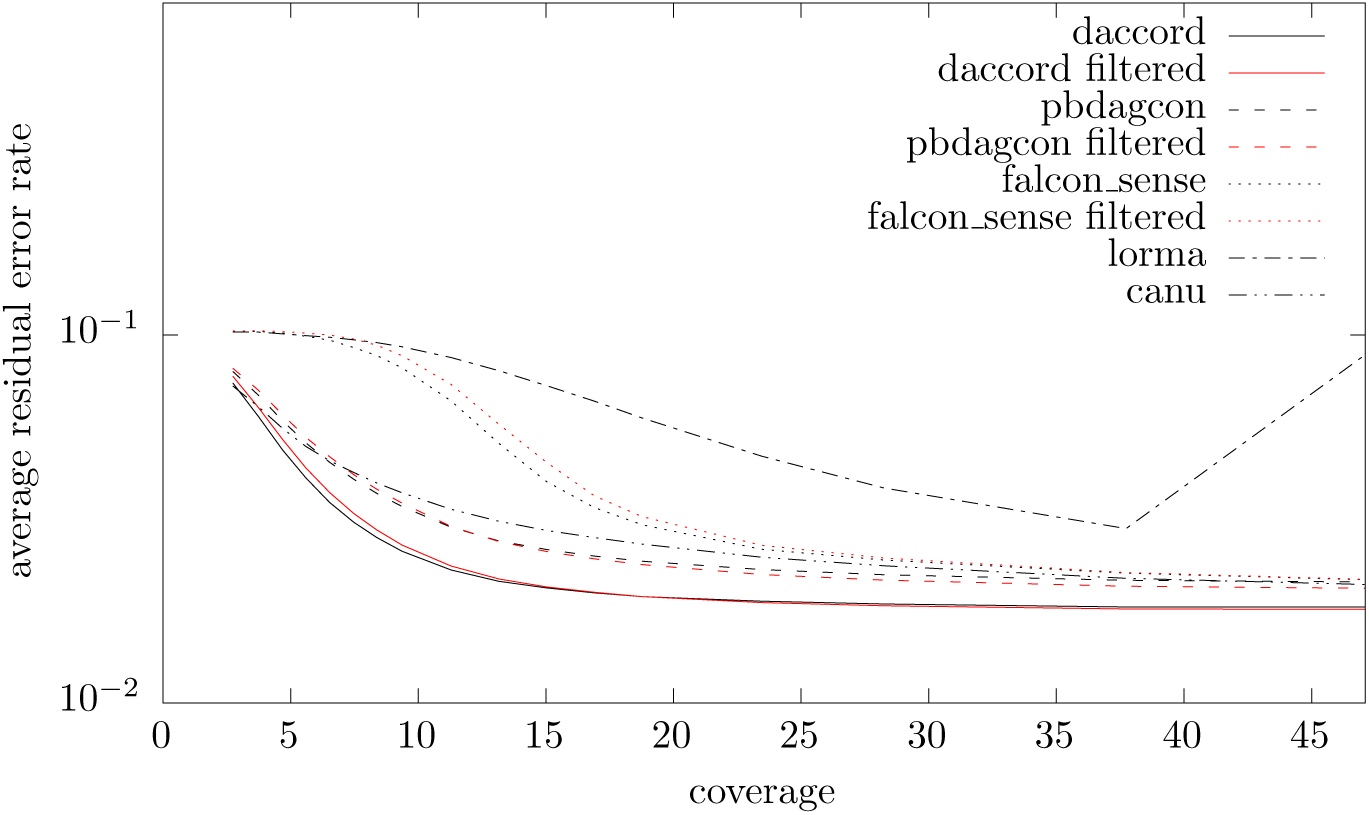
Comparison of reconstruction quality for real 2d Oxford Nanopore E. coli data

**Figure 27:**
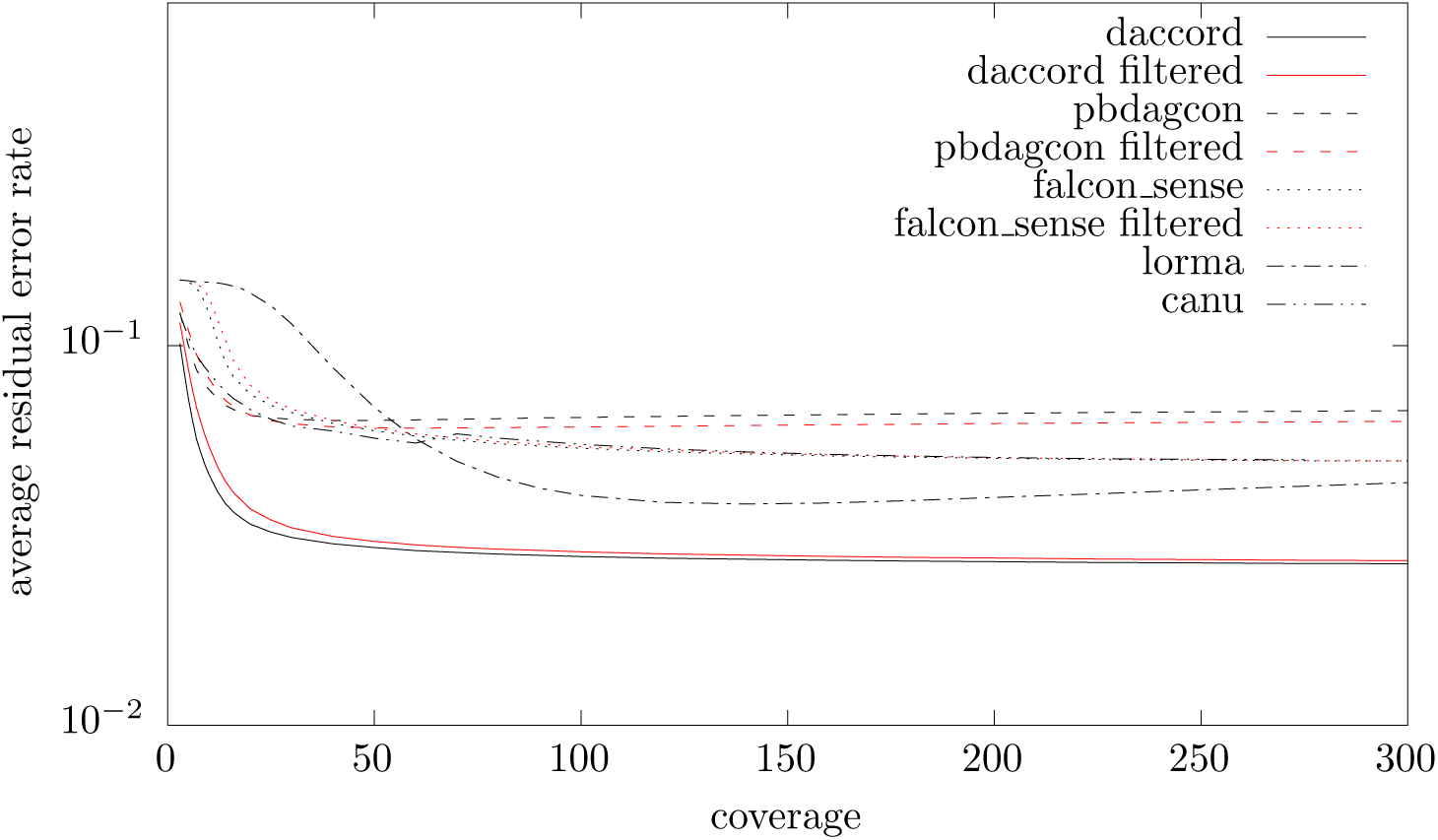
Comparison of reconstruction quality for real 1d Oxford Nanopore E. coli data

**Figure 28:**
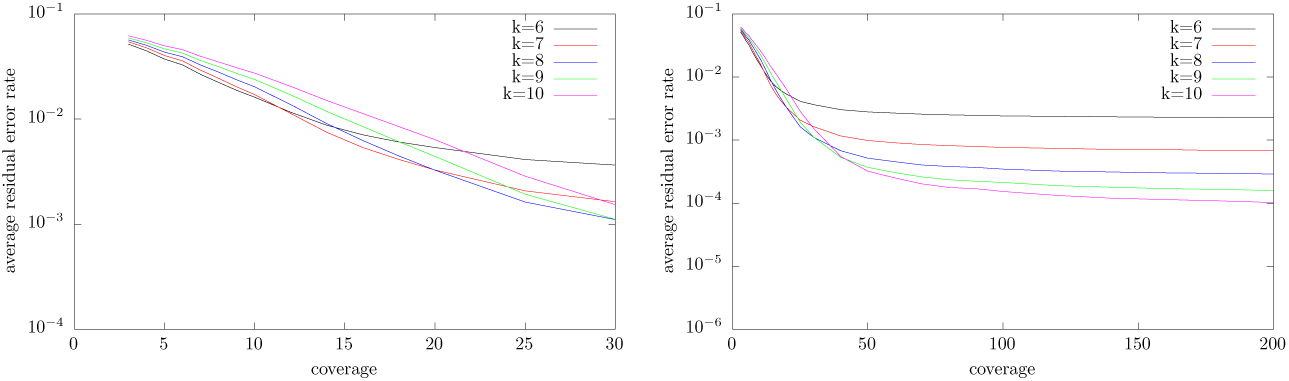
Dependence of reconstruction quality (average error rate per read) on k for k = 6; 7; 8; 9; 10 for simulated S. cerevisiae data. The left graph shows sequencing depth 3 to 30, the right graph 3 to 200. Note that the vertical axes are logarithmic.

**Figure 29:**
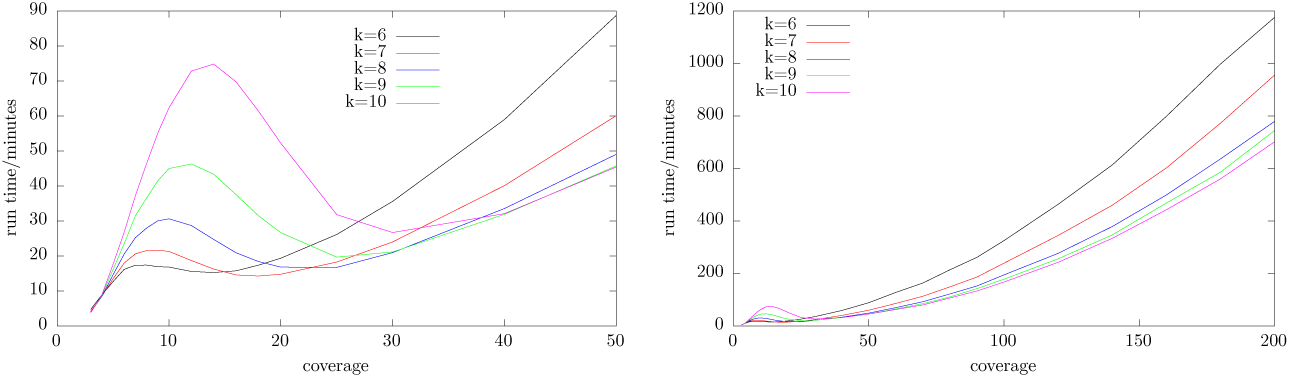
Dependence of run-time on k for k = 6; 7; 8; 9; 10 for simulated S. cerevisiae data. The left graph shows sequencing depth 3 to 50, the right graph 3 to 200.

**Table 3:**
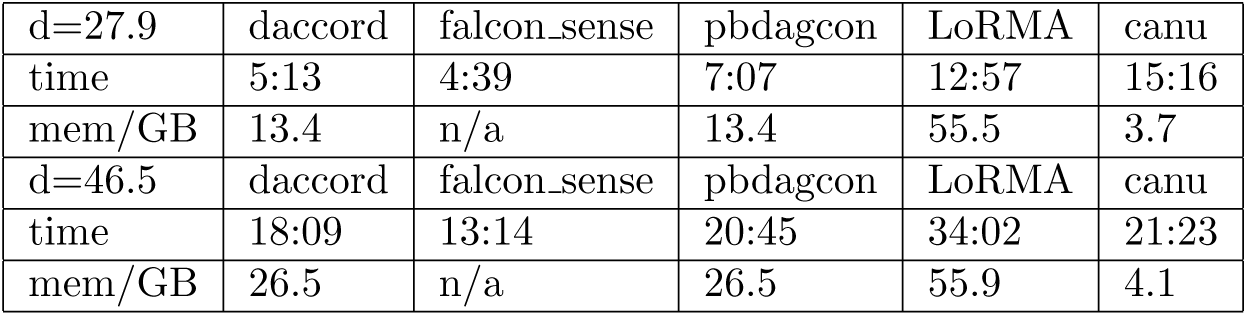
Run-time and memory usage of tools on real Oxford Nanopore sequenced 2D E. coli data

**Table 4:** Run-time and memory usage of tools on real Oxford Nanopore sequenced 1D E. coli data

|  |  |  |  |  |  |
| --- | --- | --- | --- | --- | --- |
| d=29.8 | daccord | falcon_sense | pbdagcon | LoRMA | canu |
| time | 7:13 | 5:56 | 8:48 | 6:48 | 22:53 |
| mem/GB | 9.1 | n/a | 9.1 | 55.3 | 3.8 |
| d=49.5 | daccord | falcon_sense | pbdagcon | LoRMA | canu |
| time | 18:55 | 16:19 | 24:28 | 18:17 | 29:56 |
| mem/GB | 15.8 | n/a | 15.8 | 55.7 | 4.7 |

## Conclusion

We have presented a new method for long hybrid long read error correction. The approach is practical and our implementation outperforms state of the art competitors in terms of reconstruction accuracy. In upcoming work we will discuss more involved alignment filtering to remove repeat induced alignments and thus improve error correction for long repeat regions. While our approach works for data produced by Oxford Nanopore sequencers the resulting corrected data still contains a high rate of errors. Solving this may require a more sophisticated model for sequencing events.

## A Computing Edit Scripts Between Reads

W.l.o.g. assume *P*_i_.*b* ≤ *P*_k_.*b*, the other case is symmetric. Let ‘ denote the minimum length of a prefix of *P*_i_.*S* s.t. Occ_{*M*,*D*,*S*__A,*S*__C,*S*__G,*S*__T}_(*P*_i_. *S*_1.‘_) = *P*_k_.*b* − *P*_i_.*b*. We remove the prefix *S*_1;‘_ from *P*_i_.*S* and update *P*_i_.*b* to *P*_k_.*b*. This is also ^equivalent to clipping o the rst Occ^{*M*,*I*_A_,*I*_C_,*I*_G_,*I*_T_,*S*_A_,*S*_C_,*S*_G_,*S*_T_}^(*P*^i^.*S*^1.‘^) bases^ off read(*P*_i_). If *P*_i_.*e* ≠ *P*_k_.*e* then w.l.o.g. assume *P*_i_.*e* ≤ *P*_k_.*e*. Again let *r* denote the length of a minimum length suffix of *P*_k_.*S* s.t. Occ_{*M*,*D*,*S*__A,*S*__C,*S*__G,*S*__T}_^(*P*^k^.*S*^|*P*_k_.*S*| − *r*, |*P*_k_.*S*|^)=*P*^k^.*e*−*P*^i^.*e*. We remove the suffix^ *P*k^.*S*^|*P*_k_.*S*|−*r*, |*P*_k_.*S*| from *P*_k_.*S* and update *P*_k_.*e* to *P*_i_.*e*. This is also equivalent to clipping off ^the last Occ^{*M*,*I*_A_,*I*_C_,*I*_G_,*I*_T_,*S*_A_,*S*_C_,*S*_G_,*S*_T_}^(*P*^k^.*S*^|*P*_k_.*S*|−*r*, |*P*_k_.*S*|^) bases off^ ^read(*P*^k^).^ These steps ensure [*P*_i_.*b*, *P*_i_.*e*] = [*P*_k_.*b*, *P*_k_.*e*] while keeping *P*_i_.*S* and *P*_k_.*S* ad^missible for^^*G*^*P*_i_.*j*_*P*_i_.*b*,*P*_i_.*e*_^and^^*G*^*P*_k_.*j*_*P*_k_.*b*,*P*_k_.*e*_^respectively. In particular we have Occ^{*M*, *D*,*S*_A_,*S*_C_,*S*_G_,*S*_T_}^(*P*^i^.*S*) = Occ^{*M*, *D*,*S*_A_,*S*_C_,*S*_G_,*S*_T_}^(*P*^k^.*S*) =*P*^i^.*e*−*P*^i^.*b*+ 1.^ To deduce the final edit script we parse the remainders of *P*_i_.*S* and *P*_k_.*S* from left to right. To this end let *R*_i_ = read(*P*_i_) and *R*_k_ = read(*P*_k_). We perform the following two steps *P*_i_.*e* − *P*_i_.*b*+ 1 times. First let *P*_i_.*S*_1;y_ and *P*_k_.*S*_1,z_ denote the maximum length prefixes of *P*_i_.*S* and *P*_k_.*S* respectively s.t. that all operations contained in these pre xes are insertions. W.l.o.g. let *y* ≤ *z*, the other case is symmetric. Let *C*_i_ = *R*_i1.y_ and *C*_k_ = *R*_k1.y_. Scan *C*_i_ and *C*_k_ from left to right. If we nd the same symbol in a position then append an *M* operation to the edit script. Otherwise append *S*_X_ where *X* denotes the symbol seen in *C*_k_. Let *E*_k_ = *R*_k__y+1.z_. For each symbol *X* in *E*_k_ scanned from left to right append *I*_X_ to the edit script (in the case of *y* > *z* we would add deletions instead). Now remove the prefixes *P*_i_.*S*_1,y_ and *P*_k_.*S*_1.z_ from *P*_i_ and *P*_k_ respectively and the prefixes *R*_i1,y_ and *R*_k1,z_ from *R*_i_ and *R*_k_ respectively. As the second step we are handling the next non insertion operation Y and *Z* in *P*_i_.*S* and *P*_k_.*S* respectively. If *Y* = *Z* = *D* then we add nothing to the edit script. If *Y* = *D* ≠ Z then we append the operation *I*_R__k1_ to the edit script and remove the rst symbol from *R*_k_. If *Y* ≠ *D* = *Z* then we append a *D* operation to the edit script and remove the rst symbol from *R*_i_. If *Y* ≠ *D* and *Z* ≠ *D*, then let *c* = *R*_i1_ and *d* = *R*_k1_. If *c* = *d* we append an *M* to the edit script, otherwise we append an *S*_d_. In both cases we remove the rst symbol from *R*_i_ and R_k_. To conclude the second step we remove the rst operation from *P*_i_.*S* and *P*_k_.*S* respectively. Finally there may be some insertion operations in *P*_i_.*S* or *P*_j_.*S* left. We handle these by running step one once more.

## B Figures for S288C

**Figure 30:**
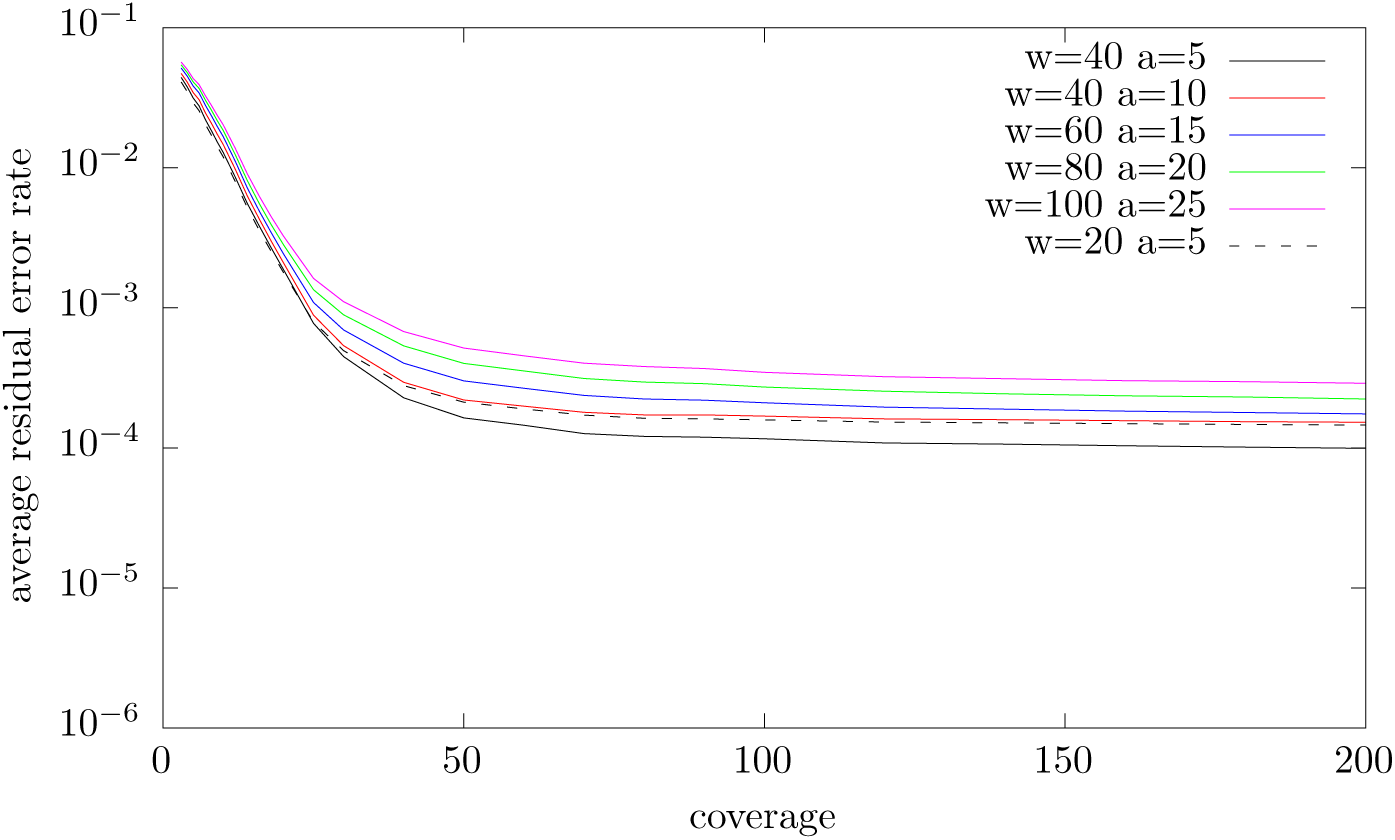
Dependence of reconstruction quality (average error rate per read) on window parameters w and a for simulated S. cerevisiae data.

**Figure 31:**
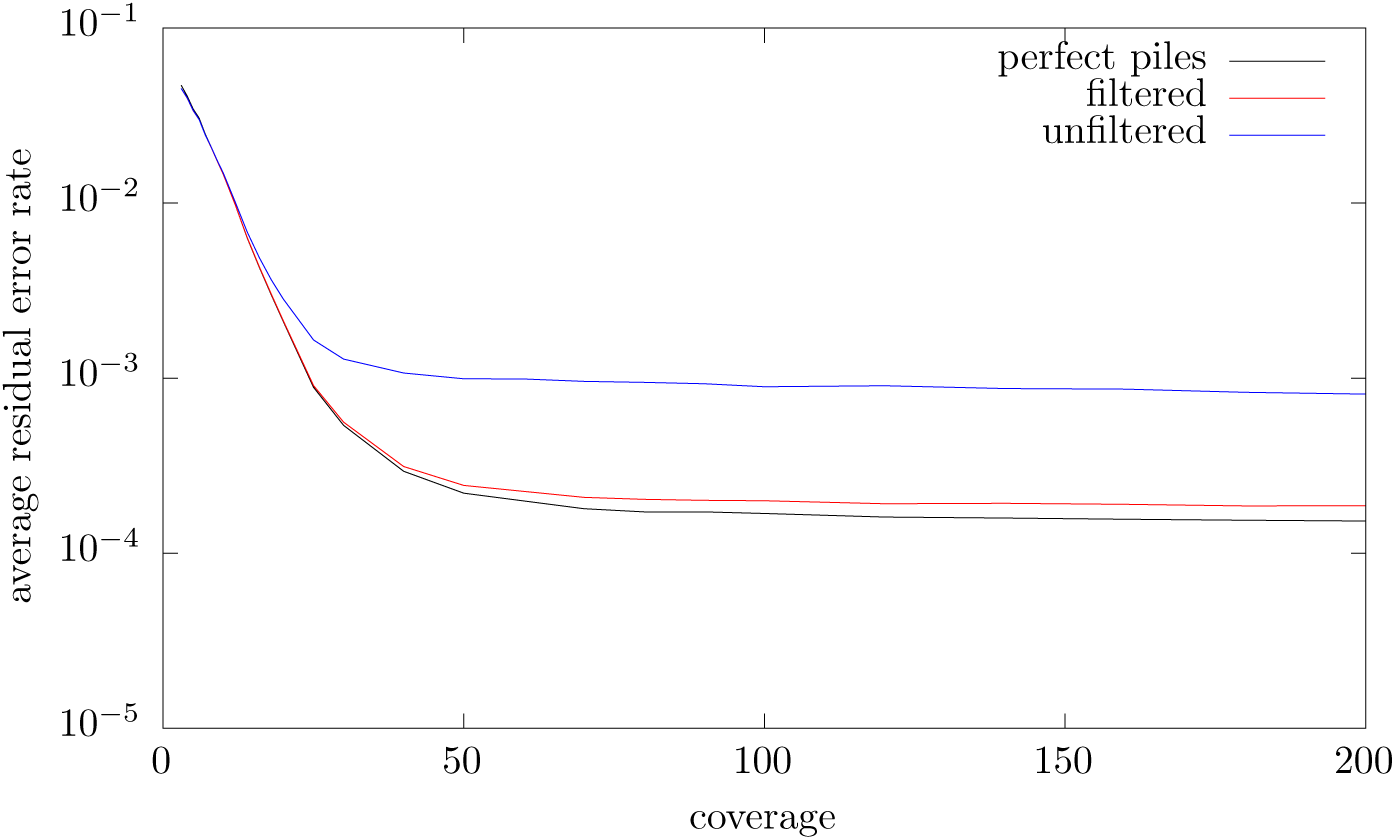
Dependence of reconstruction quality (average error rate per read) on filtering for simulated S. cerevisiae data.

**Figure 32:**
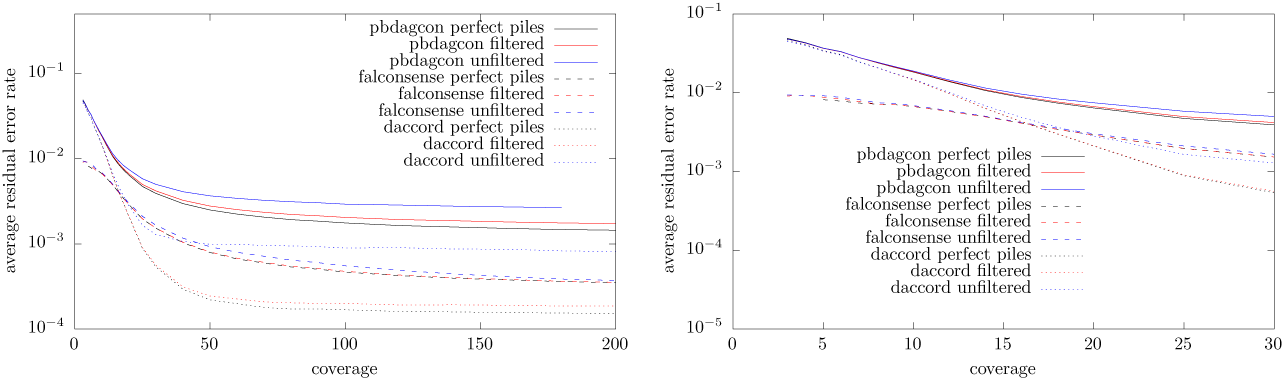
Comparison with pbdagcon falcon sense for simulated S. cerevisiae data

**Figure 33:**
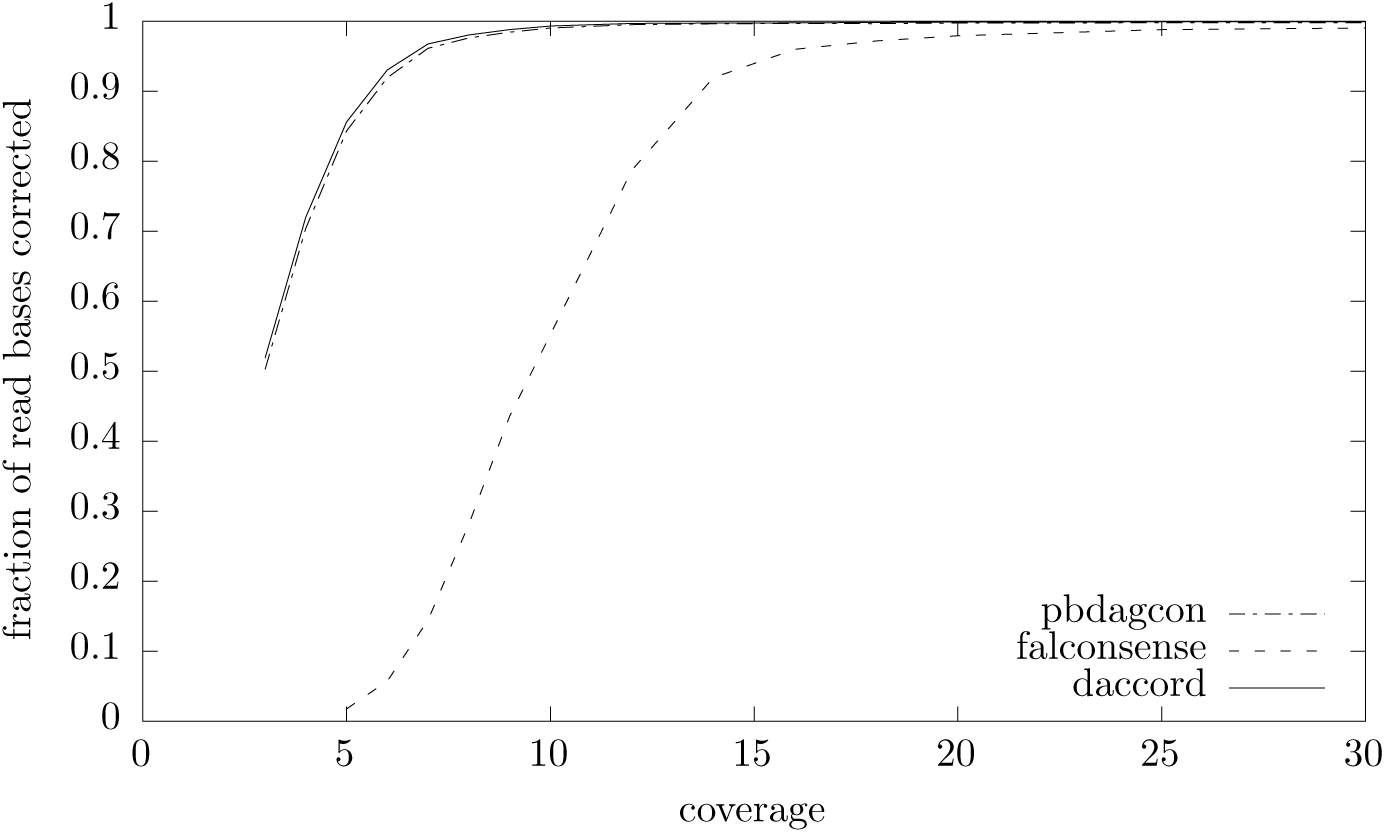
Fraction of read bases corrected by daccord, pbdagcon and falcon sense on perfect piles of simulated S. cerevisiae data

**Figure 34:**
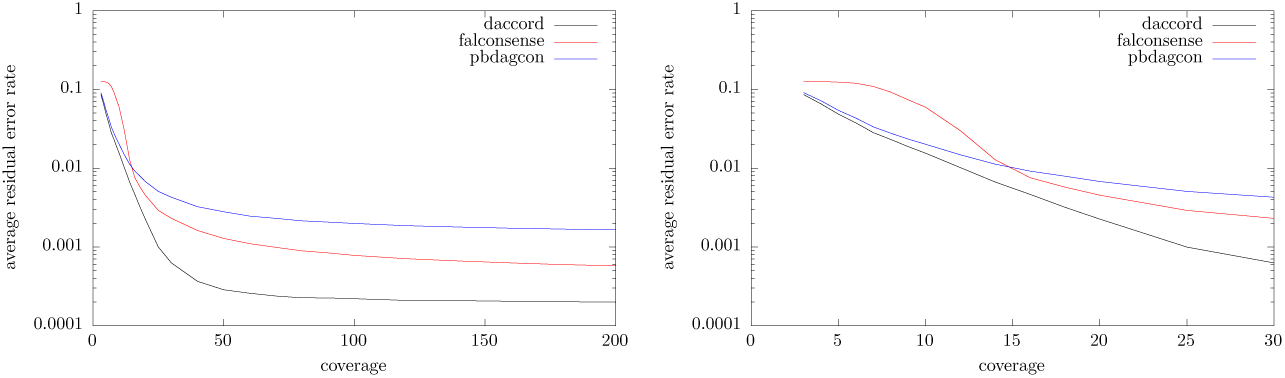
Comparison of reconstruction quality on perfect alignment piles of simulated E. coli data. Output data was inserted into the original reads for comparison.

**Figure 35:**
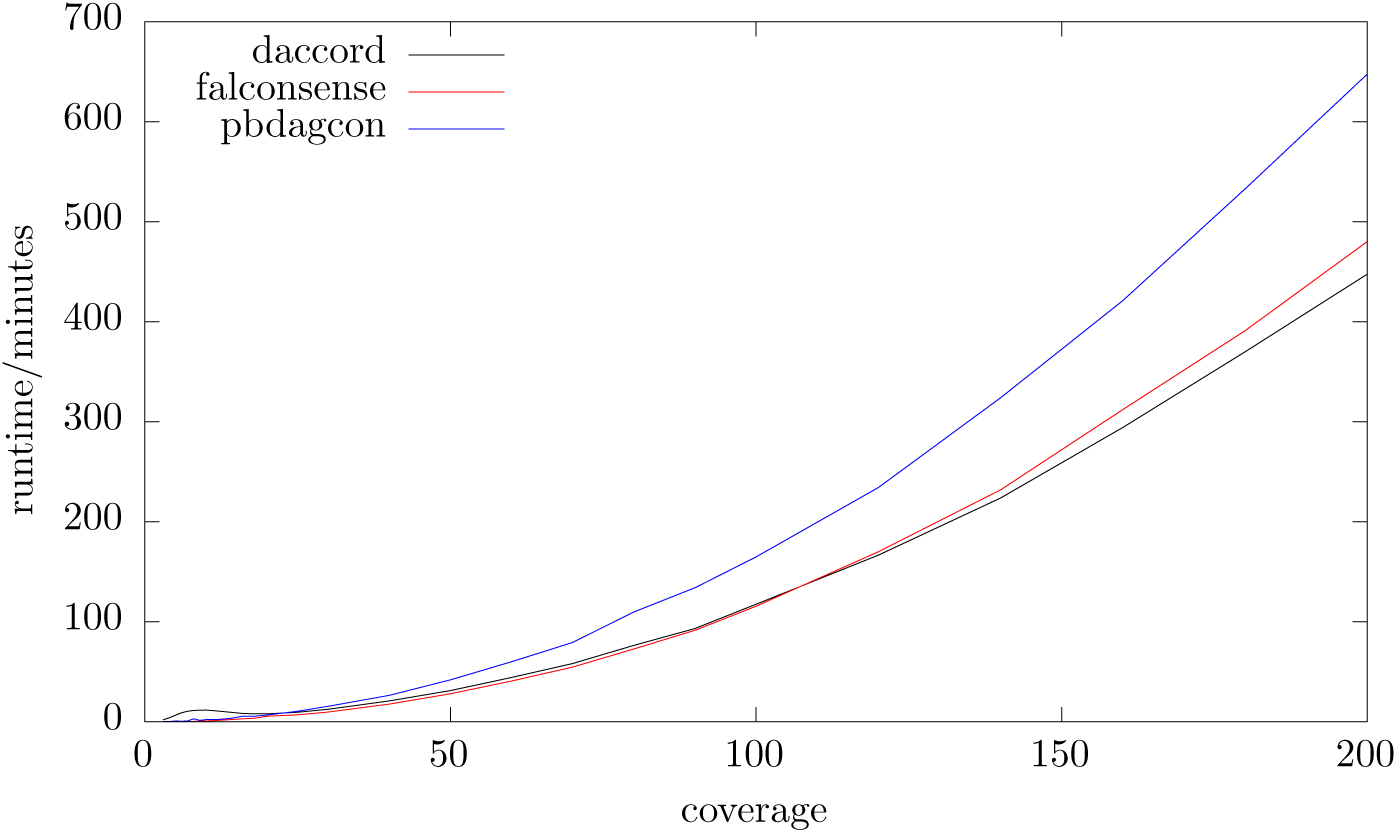
Run-time comparison with pbdagcon and falcon sense on perfect piles of simulated S. cerevisiae data

**Figure 36:**
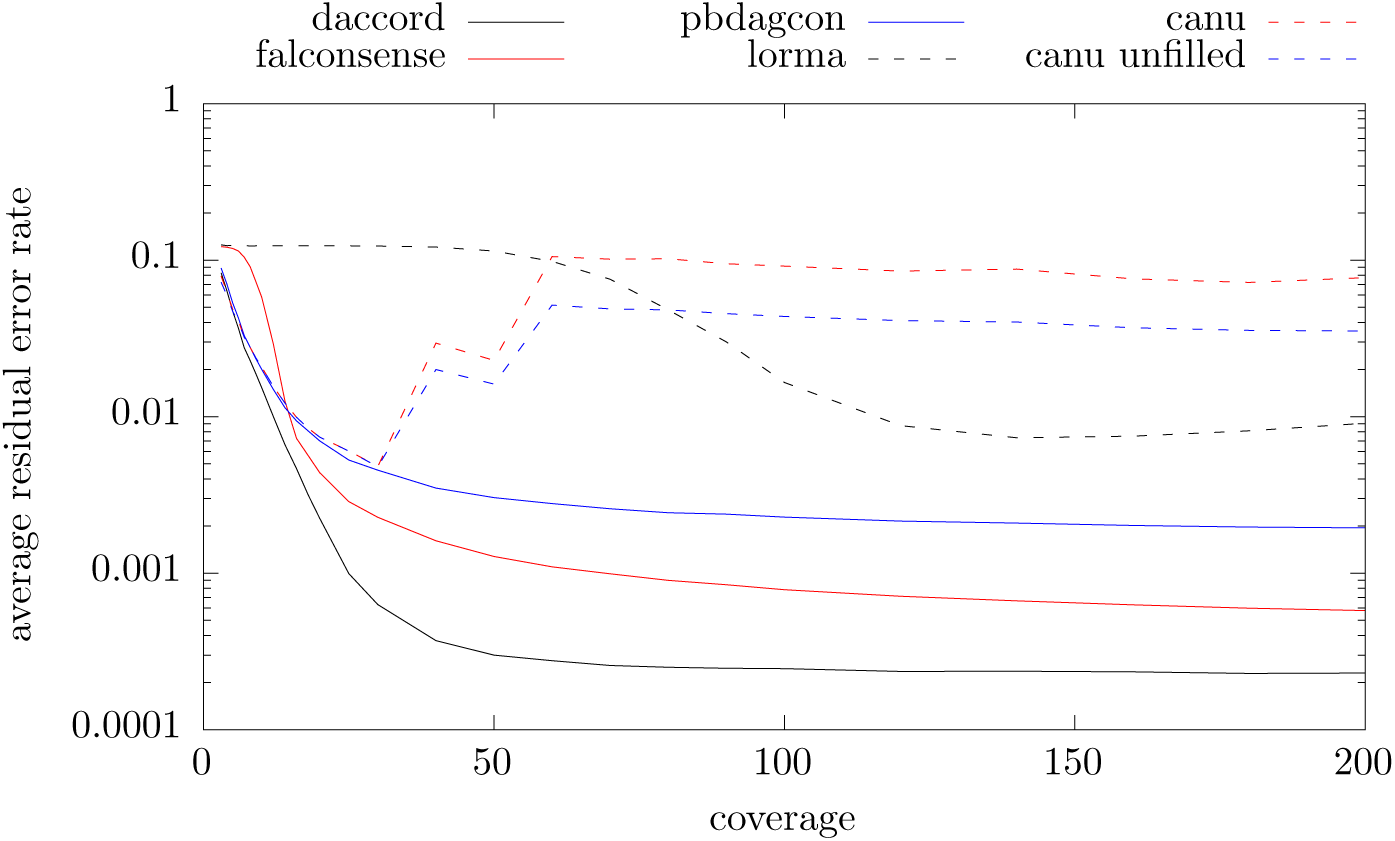
6: Comparison of correction performance with pbdagcon, falcon sense, LoRMA and Canu for simulated S. cerevisiae data

**Figure 37:**
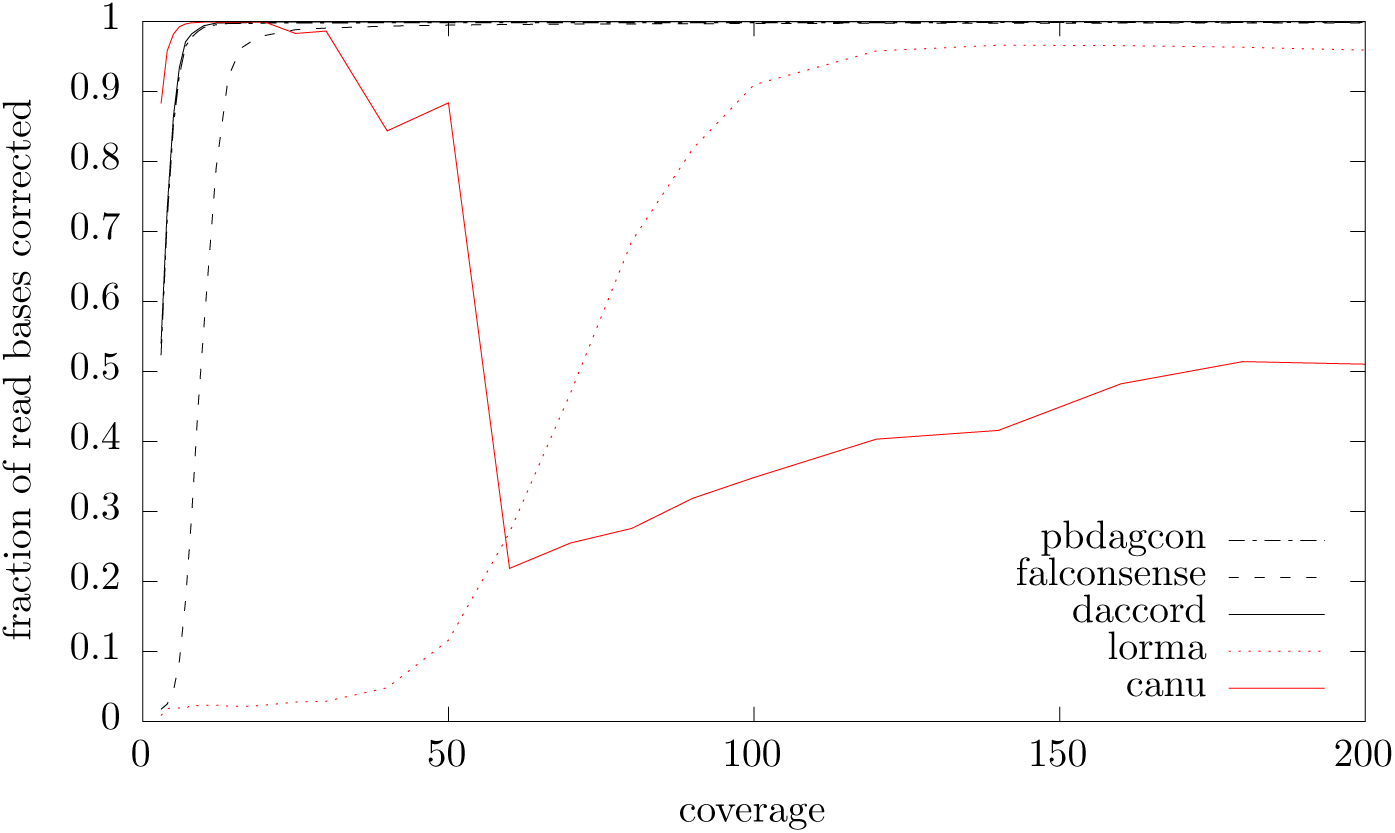
Fraction of read bases corrected by daccord, pbdagcon, falcon sense, LoRMA and Canu on simulated S. cerevisiae data

**Figure 38:**
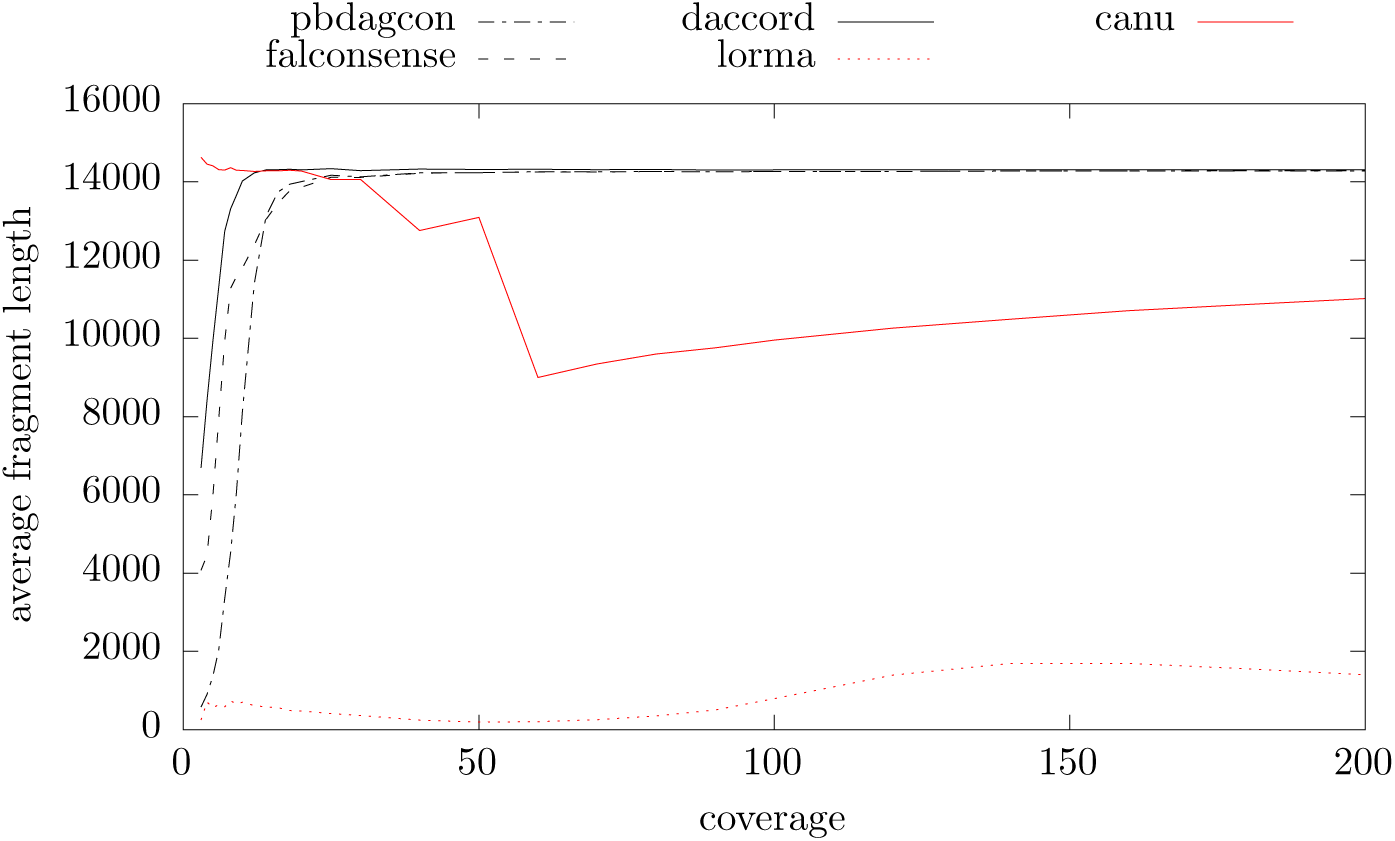
Average corrected fragment length produced by daccord, pbdagcon, falcon sense, LoRMA and Canu on simulated S. cerevisiae data

## References

1. Paci cBiosciences DevNet E. coli long read data. https://github.com/PacificBiosciences/DevNet/wiki/E-coli-Bacterial-Assembly.

2. Paci cBiosciencesDevNetSaccharomycescerevisiaelongread data. https://github.com/PacificBiosciences/DevNet/wiki/Saccharomyces-cerevisiae-W303-Assembly-Contigs.

3. M. D. Adams, S. E. Celniker, R. A. Holt, C. A. Evans, J. D. Gocayne, P. G. Amanatides, S. E. Scherer, P. W. Li, R. A. Hoskins, R. F. Galle, R. A. George, S. E. Lewis, S. Richards, M. Ashburner, S. N. Henderson, G. G. Sutton, J. R. Wortman, M. D. Yandell, Q. Zhang, L. X. Chen, R. C. Brandon, Y.-H. C. Rogers, R. G. Blazej, M. Champe, B. D. Pfei er, K. H. Wan, C. Doyle, E. G. Baxter, G. Helt, C. R. Nelson, G. L. M. Gabor, J. F. Abril, A. Agbayani, H.-J. An, C. Andrews-Pfannkoch, D. Baldwin, R. M. Ballew, M. Basu, J. Baxendale, L. Bayraktaroglu, E. M. Beasley, K. Y. Beeson, P. V. Benos, B. P. Berman, D. Bhandari, S. Bolshakov, D. Borkova, M. R. Botchan, J. Bouck, P. Brokstein, P. Brottier, K. C. Burtis, D. A. Busam, H. Butler, E. Cadieu, A. Center, I. Chandra, J. M. Cherry, S. Cawley, C. Dahlke, L. B. Davenport, P. Davies, B. d. Pablos, A. Delcher, Z. Deng, A. D. Mays, I. Dew, S. M. Dietz, K. Dodson, L. E. Doup, M. Downes, S. Dugan-Rocha, B. C. Dunkov, P. Dunn, K. J. Durbin, C. C. Evangelista, C. Ferraz, S. Ferriera, W. Fleischmann, C. Fosler, A. E. Gabrielian, N. S. Garg, W. M. Gelbart, K. Glasser, A. Glodek, F. Gong, J. H. Gorrell, Z. Gu, P. Guan, M. Harris, N. L. Harris, D. Harvey, T. J. Heiman, J. R. Hernandez, J. Houck, D. Hostin, K. A. Houston, T. J. Howland, M.-H. Wei, C. Ibegwam, M. Jalali, F. Kalush, G. H. Karpen, Z. Ke, J. A. Kennison, K. A. Ketchum, B. E. Kimmel, C. D. Kodira, C. Kraft, S. Kravitz, D. Kulp, Z. Lai, P. Lasko, Y. Lei, A. A. Levitsky, J. Li, Z. Li, Y. Liang, X. Lin, X. Liu, B. Mattei, T. C. McIntosh, M. P. McLeod, D. McPherson, G. Merkulov, N. V. Milshina, C. Mobarry, J. Morris, A. Moshrefi, S. M. Mount, M. Moy, B. Murphy, L. Murphy, D. M. Muzny, D. L. Nelson, D. R. Nelson, K. A. Nelson, K. Nixon, D. R. Nusskern, J. M. Pacleb, M. Palazzolo, G. S. Pittman, S. Pan, J. Pollard, V. Puri, M. G. Reese, K. Reinert, K. Remington, R. D. C. Saunders, F. Scheeler, H. Shen, B. C. Shue, I. Siden-Kiamos, M. Simpson, M. P. Skupski, T. Smith, E. Spier, A. C. Spradling, M. Stapleton, R. Strong, E. Sun, R. Svirskas, C. Tector, R. Turner, E. Venter, A. H. Wang, X. Wang, Z.-Y. Wang, D. A. Wassarman, G. M. Weinstock, J. Weissenbach, S. M. Williams, T. Woodage, K. C. Worley, D. Wu, S. Yang, Q. A. Yao, J. Ye, R.-F. Yeh, J. S. Zaveri, M. Zhan, G. Zhang, Q. Zhao, L. Zheng, X. H. Zheng, F. N. Zhong, W. Zhong, X. Zhou, S. Zhu, X. Zhu, H. O. Smith, R. A. Gibbs, E. W. Myers, G. M. Rubin, and J. C. Venter. The genome sequence of drosophila melanogaster. Science, 287(5461):2185-2195, 2000.

4. K. F. Au, J. G. Underwood, L. Lee, and W. H. Wong. Improving pacbio long-read accuracy by short read alignment. PLOS ONE, 7(10):1–8, 10 2012.

5. K. Berlin, S. Koren, C.-S. Chin, J. P. Drake, J. M. Landolin, and A. M. Phillippy. Assembling large genomes with single-molecule sequencing and locality-sensitive hashing. Nat Biotech, 33(6):623–630, Jun 2015. Research.

6. C.-S. Chin, D. H. Alexander, P. Marks, A. A. Klammer, J. Drake, C. Heiner, A. Clum, A. Copeland, J. Huddleston, E. E. Eichler, S. W. Turner, and J. Korlach. Nonhybrid, finished microbial genome assemblies from long-read smrt sequencing data. Nat Meth, 10(6):563–569, Jun 2013.

7. C.-S. Chin, P. Peluso, F. J. Sedlazeck, M. Nattestad, G. T. Concepcion, A. Clum, C. Dunn, R. O’Malley, R. Figueroa-Balderas, A. Morales-Cruz, G. R. Cramer, M. Delledonne, C. Luo, J. R. Ecker, D. Cantu, D. R. Rank, and M. C. Schatz. Phased diploid genome assembly with single-molecule real-time sequencing. Nat Meth, 13(12):1050–1054, Dec 2016.

8. J. Fischer. Optimal succinctness for range minimum queries., editor, Proceedings LATIN 2010, volume 6034 of LNCS, pages 158–169. Springer, 2010.

9. T. Gagie, S. J. Puglisi, and A. Turpin. Range quantile queries: Another virtue of wavelet trees. editors, String Processing and Information Retrieval, 16th International Symposium, SPIRE 2009, Saariselka, Finland, August 25-27, 2009, Proceedings, volume. 5721 of Lecture Notes in Computer Science, pages 1–6. Springer, 2009.

10. T. Hackl, R. Hedrich, J. Schultz, and F. Forster. proovread: large-scale high-accuracy pacbio correction through iterative short read consensus. Bioinformatics, 30(21):3004–3011, Nov 2014. 25015988[pmid].

11. R. M. Idury and M. S. Waterman. A New Algorithm for DNA Sequence Assembly. Journal of Computational Biology, 2(2):291–306 Jan 1995.

12. S. Koren, G. P. Harhay, T. P. Smith, J. L. Bono, D. M. Harhay, S. D. Mcvey, D. Radune, N. H. Bergman, and A. M. Phillippy. Reducing assembly complexity of microbial genomes with single-molecule sequencing. Genome Biology, 14(9):R101, 2013.

13. S. Koren, B. P. Walenz, K. Berlin, J. R. Miller, and A. M. Phillippy. Canu: scalable and accurate long-read assembly via adaptive k-mer weighting and repeat separation. bioRxiv, 2016.

14. E. S. Lander et al. Initial sequencing and analysis of the human genome. Nature, 409(6822):860–921, Feb 2001.

15. Y. Lin, J. Yuan, M. Kolmogorov, M. W. Shen, and P. A. Pevzner. Assembly of Long Error-Prone Reads Using de Bruijn Graphs. bioRxiv, 2016.

16. N. Loman. Nanopore r9 rapid run data release. http://lab.loman.net/2016/07/30/nanopore-r9-data-release/.

17. R. Luo, B. Liu, Y. Xie, Z. Li, W. Huang, J. Yuan, G. He, Y. Chen, Q. Pan, Y. Liu, J. Tang, G. Wu, H. Zhang, Y. Shi, Y. Liu, C. Yu, B. Wang, Y. Lu, C. Han, D. W. Cheung, S.-M. Yiu, S. Peng, Z. Xiaoqian, G. Liu, X. Liao, Y. Li, H. Yang, J. Wang, T.-W. Lam, and J. Wang. Soapdenovo2: an empirically improved memory-efficient short-read de novo assembler. GigaScience, 1(1):18, 2012.

18. G. Miclotte, M. Heydari, P. Demeester, S. Rombauts, Y. Van de Peer, P. Audenaert, and J. Fostier. Jabba: hybrid error correction for long sequencing reads. Algorithms for Molecular Biology, 11(1):10, 2016.

19. E. W. Myers. Mapping your reads: damapper. https://dazzlerblog.wordpress.com/2016/07/31/damapper-mapping-your-reads/.

20. E. W. Myers. An O(N D) difference algorithm and its variations. Algorithmica, 1(1):251-266, 1986.

21. G. Myers. Efficient local alignment discovery amongst noisy long reads., editors, Algorithms in Bioinformatics - 14th International Workshop, WABI 2014, Wroclaw, Poland, September 8-10, 2014. Proceedings, volume 8701 of Lecture Notes in Computer Science, pages 52–67. Springer, 2014.

22. Y. Ono, K. Asai, and M. Hamada. PBSIM: pacbio reads simulator - toward accurate genome assembly. Bioinformatics, 29(1):119–121, 2013.

23. P. A. Pevzner, H. Tang, and M. S. Waterman. An Eulerian path approach to DNA fragment assembly. Proceedings of the National Academy of Sciences, 98(17):9748–9753, 2001.

24. L. Salmela and E. Rivals. Lordec: accurate and efficient long read error correction. Bioinformatics, 30(24):3506, 2014.

25. L. Salmela, R. Walve, E. Rivals, and E. Ukkonen. Accurate self-correction of errors in long reads using de bruijn graphs. Bioinformatics, 2016.

26. J. T. Simpson, K. Wong, S. D. Jackman, J. E. Schein, S. J. M. Jones, and Birol. ABySS: A parallel assembler for short read sequence data. Genome Research, 19(6):1117–1123, June 2009.

27. G. Tischler. Benchmarking damapper. https://dazzlerblog.wordpress.com/2016/08/24/benchmarking-damapper/.

28. J. C. Venter, M. D. Adams, E. W. Myers, P. W. Li, R. J. Mural, G. G. Sutton, H. O. Smith, M. Yandell, C. A. Evans, R. A. Holt, J. D. Gocayne, P. Amanatides, R. M. Ballew, D. H. Huson, J. R. Wortman, Q. Zhang, C. D. Kodira, X. H. Zheng, L. Chen, M. Skupski, G. Subramanian, P. D. Thomas, J. Zhang, G. L. Gabor Miklos, C. Nelson, S. Broder, A. G. Clark, J. Nadeau, V. A. McKusick, N. Zinder, A. J. Levine, R. J. Roberts, M. Simon, C. Slayman, M. Hunkapiller, R. Bolanos, A. Delcher, I. Dew, D. Fasulo, M. Flanigan, L. Florea, A. Halpern, S. Hannenhalli, S. Kravitz, S. Levy, C. Mobarry, K. Reinert, K. Remington, J. Abu-Threideh, E. Beasley, K. Biddick, V. Bonazzi, R. Brandon, M. Cargill, I. Chandramouliswaran, R. Charlab, K. Chaturvedi, Z. Deng, V. D. Francesco, P. Dunn, K. Eilbeck, C. Evangelista, A. E. Gabrielian, W. Gan, W. Ge, F. Gong, Z. Gu, P. Guan, T. J. Heiman, M. E. Higgins, R.-R. Ji, Z. Ke, K. A. Ketchum, Z. Lai, Y. Lei, Z. Li, J. Li, Y. Liang, X. Lin, F. Lu, G. V. Merkulov, N. Milshina, H. M. Moore, A. K. Naik, V. A. Narayan, B. Neelam, D. Nusskern, D. B. Rusch, S. Salzberg, W. Shao, B. Shue, J. Sun, Z. Y. Wang, A. Wang, X. Wang, J. Wang, M.-H. Wei, R. Wides, C. Xiao, C. Yan, A. Yao, J. Ye, M. Zhan, W. Zhang, H. Zhang, Q. Zhao, L. Zheng, F. Zhong, W. Zhong, S. C. Zhu, S. Zhao, D. Gilbert, S. Baumhueter, G. Spier, C. Carter, A. Cravchik, S. Woodage, F. Ali, H. An, A. Awe, D. Baldwin, H. Baden, M. Barnstead, I. Barrow, K. Beeson, D. Busam, A. Carver, A. Center, M. L. Cheng, L. Curry, S. Danaher, L. Davenport, R. Desilets, S. Dietz, K. Dodson, L. Doup, S. Ferriera, N. Garg, A. Gluecksmann, B. Hart, J. Haynes, C. Haynes, C. Heiner, S. Hladun, D. Hostin, J. Houck, T. Howland, C. Ibegwam, J. Johnson, F. Kalush, L. Kline, S. Koduru, A. Love, F. Mann, C. May, S. McCawley, T. McIntosh, I. McMullen, M. Moy, L. Moy, B. Murphy, K. Nelson, C. Pfannkoch, E. Pratts, V. Puri, H. Qureshi, M. Reardon, R. Rodriguez, Y.-H. Rogers, D. Romblad, B. Ruhfel, R. Scott, C. Sitter, M. Smallwood, E. Stewart, R. Strong, E. Suh, R. Thomas, N. N. Tint, R. Tse, C. Vech, G. Wang, J. Wetter, S. Williams, M. Williams, S. Windsor, E. Winn-Deen, K. Wolfe, J. Zaveri, K. Zaveri, J. F. Abril, R. Guigo, M. J. Campbell, K. V. Sjolander, B. Karlak, A. Kejariwal, H. Mi, B. Lazareva, T. Hatton, A. Narechania, K. Diemer, A. Muruganujan, N. Guo, S. Sato, V. Bafna, S. Istrail, R. Lippert, R. Schwartz, B. Walenz, S. Yooseph, D. Allen, A. Basu, J. Baxendale, L. Blick, M. Caminha, J. Carnes-Stine, P. Caulk, Y.-H. Chiang, M. Coyne, C. Dahlke, A. D. Mays, M. Dombroski, M. Donnelly, D. Ely, S. Esparham, C. Fosler, H. Gire, S. Glanowski, K. Glasser, A. Glodek, M. Gorokhov, K. Graham, B. Gropman, M. Harris, J. Heil, S. Henderson, J. Hoover, D. Jennings, C. Jordan, J. Jordan, J. Kasha, L. Kagan, C. Kraft, A. Levitsky, M. Lewis, X. Liu, J. Lopez, D. Ma, W. Majoros, J. McDaniel, S. Murphy, M. Newman, T. Nguyen, N. Nguyen, M. Nodell, S. Pan, J. Peck, M. Peterson, W. Rowe, R. Sanders, J. Scott, M. Simpson, T. Smith, A. Sprague, T. Stockwell, R. Turner, E. Venter, M. Wang, M. Wen, D. Wu, M. Wu, A. Xia, A. Zandieh, and X. Zhu. The sequence of the human genome. Science, 291(5507):1304–1351, 2001.

